# The host RNA polymerase II C-terminal domain is the anchor for replication of the influenza virus genome

**DOI:** 10.1101/2023.08.01.551436

**Authors:** T Krischuns, B Arragain, C Isel, S Paisant, M Budt, T Wolff, S Cusack, N Naffakh

**Affiliations:** RNA Biology of Influenza Virus, Institut Pasteur, Université Paris Cité, CNRS UMR 3569, Paris, France; European Molecular Biology Laboratory, Grenoble, France; Unit 17 “Influenza and other Respiratory Viruses”, Robert Koch Institut, Berlin, Germany

**Keywords:** Influenza virus, RNA-dependent RNA polymerase, cellular RNA polymerase II, ANP32, transcription, replication

## Abstract

The current model is that the influenza virus polymerase (FluPol) binds either to host RNA polymerase II (RNAP II) or to the acidic nuclear phosphoprotein 32 (ANP32), which drives its conformation and activity towards transcription or replication of the viral genome, respectively. Here, we provide evidence that the FluPol-RNAP II binding interface has a so far overlooked function for replication of the viral genome. Using a combination of cell-based and *in vitro* approaches, we show that the RNAP II C-terminal-domain, jointly with ANP32, enhances FluPol replication activity and we propose a model in which the host RNAP II is the anchor for transcription and replication of the viral genome. Our data open new perspectives on the spatial coupling of viral transcription and replication and the coordinated balance between these two activities.

## Introduction

Influenza viruses are major human pathogens which cause annual epidemics (type A, B and C viruses) (Krammer et al., 2018) and have zoonotic and pandemic potential (type A viruses) (Harrington et al., 2021). The viral RNA dependent RNA polymerase (FluPol) is a key determinant of viral host-range and pathogenicity, and a prime target for antiviral drugs (Mifsud et al., 2019). It is a ∼260 kDa heterotrimer composed of the subunits PB1 (polymerase basic protein 1), PB2 (polymerase basic protein 2) and PA (polymerase acidic protein) (Wandzik et al., 2021). In viral particles each of the eight negative-sense RNA genomic segments (vRNAs) is associated with one copy of the viral polymerase and encapsidated with multiple copies of the nucleoprotein (NP) to form viral ribonucleoproteins (vRNPs) (Eisfeld et al., 2015). Upon viral infection, incoming vRNPs are imported into the nucleus in which FluPol performs transcription and replication of the viral genome through distinct primed (Plotch et al., 1979) and unprimed (Deng et al., 2006) RNA synthesis mechanisms, respectively. Determination of high-resolution FluPol structures by X-ray crystallography and cryogenic electron microscopy (cryo-EM) have revealed that a remarkable conformational flexibility allows FluPol to perform mRNA transcription (Dias et al., 2009; Guilligay et al., 2008; Kouba et al., 2019; Pflug et al., 2018; Wandzik et al., 2020). Recently, first insights into the molecular process of FluPol genome replication were revealed, however the molecular details remain largely to be determined (Carrique et al., 2020; Keown et al., 2022; Wang et al., 2022).

Transcription of negative-sense vRNAs into mRNAs occurs via a process referred to as “cap-snatching”, whereby nascent 5’-capped host RNA polymerase II (RNAP II) transcripts are bound by the PB2 cap-binding domain (Guilligay et al., 2008), cleaved by the PA endonuclease domain (Dias et al., 2009), and used as capped primers to initiate transcription (Kouba et al., 2019). Polyadenylation is achieved by a noncanonical stuttering mechanism at a 5′-proximal oligo(U) polyadenylation signal present on each vRNA (Poon et al., 1999; Wandzik et al., 2020). Influenza mRNAs therefore harbour 5’-cap and 3’-poly(A) structures characteristic of cellular mRNAs and are competent for translation by the cellular machinery. The accumulation of primary transcripts of incoming vRNPs leads to *de novo* synthesis of viral proteins, including FluPol and NP, which are thought to trigger genome replication upon nuclear import (Zhu et al., 2023).

Replication generates exact, full-length complementary RNAs (cRNAs) which in turn serve as templates for the synthesis of vRNAs. Unlike transcription, replication occurs through a primer-independent, *de novo* initiation mechanism which occurs either at the first nucleotide of the 3’-vRNA template or at nucleotide 4 and 5 of the 3’-cRNA template followed by realignment prior to elongation (Deng et al., 2006; Oymans and Te Velthuis, 2018; Zhang et al., 2010). Nascent cRNAs and vRNAs are encapsidated by NP and FluPol to form cRNPs and vRNPs, respectively (York et al., 2013). Progeny vRNPs can then perform secondary transcription and replication or, late during the infection cycle, be exported from the nucleus to the cytoplasm to be incorporated into new virions.

FluPol associates dynamically with many cellular proteins, among which the interaction with the host RNAP II transcription machinery (Krischuns et al., 2021) and the nuclear acidic nuclear phosphoprotein 32 (ANP32) (Long et al., 2016) are essential for a productive viral infection. A direct interaction between FluPol and the disordered C-terminal domain (CTD) of the largest RNAP II subunit is essential for “cap-snatching” (Engelhardt et al., 2005). In mammals, the CTD consists of 52 repeats of the consensus sequence T_1_-S_2_-P_3_-T_4_-S_5_-P_6_-S_7_ (Harlen and Churchman, 2017). To perform “cap-snatching”, FluPol binds specifically to the CTD when phosphorylated on serine 5 residues (pS5 CTD), which is a hallmark of RNAP II in a paused elongation state (Core and Adelman, 2019). Structural studies revealed bipartite FluPol CTD-binding sites for influenza type A, B and C viruses (Krischuns et al., 2022; Lukarska et al., 2017; Serna Martin et al., 2018). For type A viruses, both CTD-binding sites, denoted site 1A and site 2A, are on the PA-C-terminal domain (PA-Cter) and consist of highly conserved basic residues that directly interact with pS5 moieties of the CTD (Lukarska et al., 2017).

The host ANP32 proteins consist of a structured N-terminal leucine-rich repeat (LRR) domain and a disordered C-terminal low complexity acidic region (LCAR) and are essential for influenza virus genome replication (Nilsson-Payant et al., 2022; Swann et al., 2023). Differences in avian and mammalian ANP32 proteins represent a major driver of FluPol adaptation upon zoonotic infections of mammalian species with avian influenza viruses (Baker et al., 2018; Long et al., 2016; Swann et al., 2023). A type C virus FluPol (Flu_C_Pol) co-structure with ANP32A shows that the ANP32A LRR domain binds to an asymmetrical influenza polymerase dimer, which is presumed to represent the FluPol replication complex (Carrique et al., 2020). The two FluPol conformations observed in this dimer would represent the RNA-bound replicase (FluPol_(R)_), which synthesises *de novo* genomic RNA and the encapsidase (FluPol_(E)_), which binds the outgoing replication product and nucleates formation of the progeny RNP (Zhu et al., 2023). In addition, the ANP32 LCAR domain was shown to interact with NP, leading to a model in which the ANP32 LCAR domain recruits NP to nascent exiting RNA, thus, in combination with the FluPol_(E)_, ensuring efficient co-replicatory encapsidation of *de novo* synthesised genomic RNAs (Wang et al., 2022).

Structural, biochemical and functional studies to date have led to a model in which the FluPol binds either to the pS5 CTD or to ANP32, which drives RNPs towards transcription or replication, respectively (Zhu et al., 2023). Here, we provide genetic evidence that the FluPol CTD-binding interface has a so far overlooked function in that it is essential not only for transcription but also for replication of the viral genome. We show that the CTD, jointly with ANP32, enhances FluPol replication activity and we demonstrate in structural studies that CTD-binding to the FluPol is consistent with replication activity. We therefore propose a model in which transcription and replication of the influenza virus genome occur in association to the RNAP II CTD thereby allowing an efficient switching between the two activities.

## Results

### The FluPol CTD-binding interface is essential for replication of the viral genome

Previous studies demonstrated that the FluPol PA-Cter domain is crucial for FluPol transcription by mediating the interaction between the FluPol transcriptase (FluPol_(T)_) and the host RNAP II pS5 CTD for “cap-snatching” through direct protein-protein interaction (**Fig. 1A**, FluPol_(T)_) (Krischuns et al., 2022; Lukarska et al., 2017). Recent cryo-EM structures of Flu_C_Pol suggest that the PA-Cter domain is also involved in the formation of the FluPol replication complex (Carrique et al., 2020). Based on these Flu_C_Pol structures, we generated a model of the Flu_A_Pol replicase-encapsidase complex (**Fig. 1B** FluPol_(R)_ and FluPol_(E)_). According to our model, the residues essential for FluPol binding to the CTD are surface exposed and potentially competent for CTD-binding in the FluPol_(R)_ as well as the FluPol_(E)_ conformation. Strikingly, FluPol_(E)_ CTD-binding residues are in close proximity to the ANP32-binding interface (**Fig. 1B and S1A**). We therefore hypothesised that the FluPol CTD-binding interface may have a role beyond the described association with the pS5 CTD in the FluPol_(T)_ conformation namely to facilitate assembly of the FluPol_(R)_-ANP32-FluPol_(E)_ replication complex.

**Figure 1:**
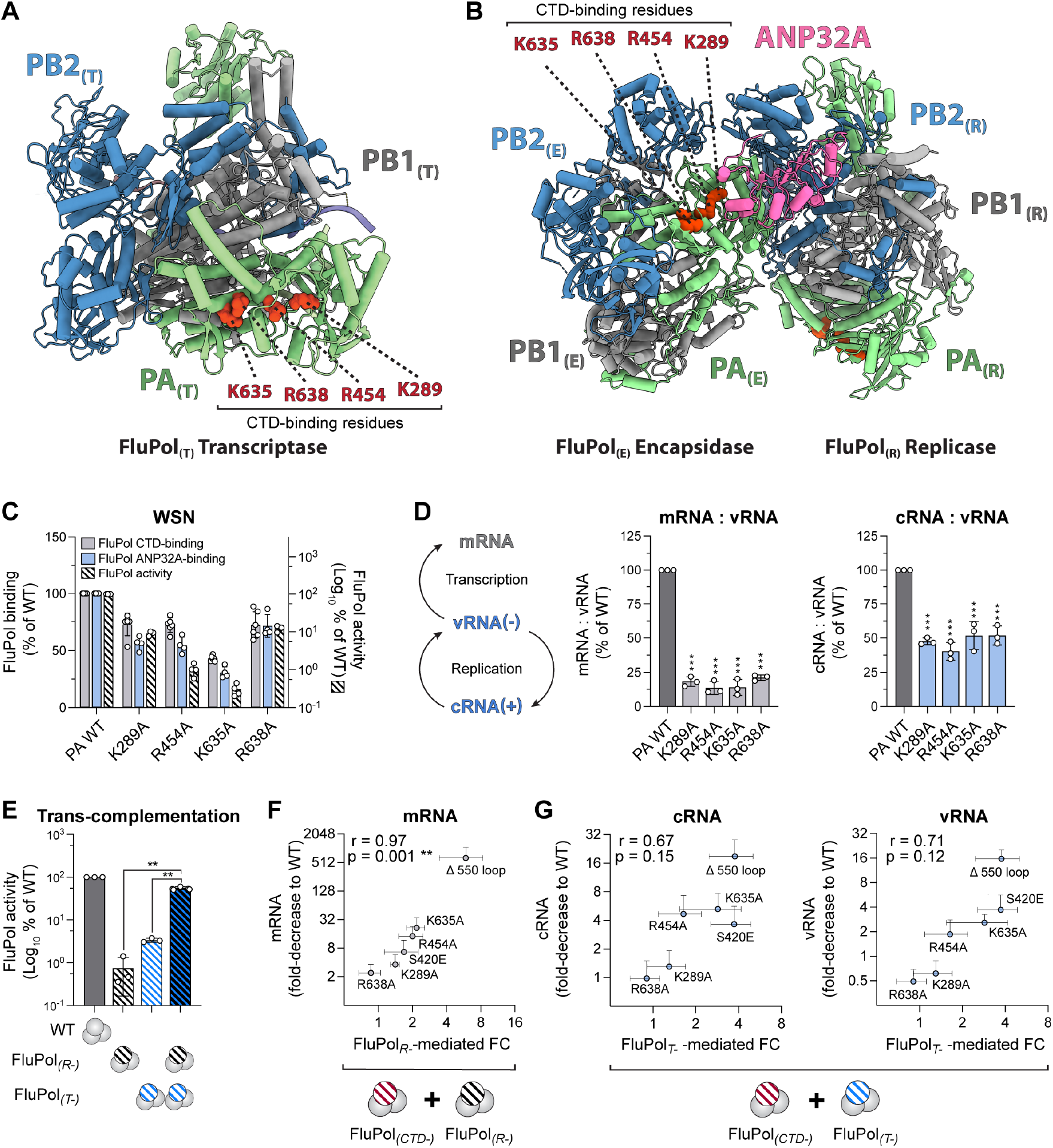
The FluPol CTD-binding interface is essential for replication of the viral genome. **(A)** Representation of the Cryo-EM structure of the Flu_A_Pol heterotrimer in the FluPol_(T)_ conformation, bound to a 3’5’ vRNA promoter and a short capped RNA primer (A/NT/60/1968, PDB: 6RR7, (Fan et al., 2019)). Ribbon diagram representation with PA in green, PB1 in grey, and PB2 in blue. Key FluPol CTD-binding residues PA K289, R454, K635 and R638 are highlighted in red. **(B)** Model of the actively replicating Zhejiang-H7N9 Flu_A_Pol replicase-ANP32-encapsidase (FluPol_(R)_, FluPol_(E)_ and ANP32A) The colour code is as in (A) with ANP32A in pink. The model was constructed by superposing equivalent Zhejiang-H7N9 Flu_A_Pol domains (elongating H7N9 Flu_A_Pol, including product-template duplex, PDB: 7QTL, (Kouba et al., 2023)) for the FluPol_(R)_ and 5’ hook bound apo-dimer Zhejiang-H7N9 Flu_A_Pol (PDB: 7ZPL, (Kouba et al., 2023)) for the FluPol_(E)_, on those of Flu_C_Pol replication complex (PDB: 6XZQ, (Carrique et al., 2020)). The ANP32A was left unchanged. Key FluPol CTD-binding residues PA K289, R454, K635 and R638 are highlighted in red on the FluPol_(R)_ as well as the FluPol_(E)._ **(C)** Cell-based WSN FluPol binding and activity assays. Left Y-axis (linear scale): FluPol binding to the CTD (grey bars) and ANP32A (blue bars) was assessed using split-luciferase-based complementation assays. HEK-293T cells were co-transfected with expression plasmids for the CTD or ANP32 tagged with one fragment of the *G. princeps* luciferase and FluPol tagged with the other fragment either with PA WT or with the indicated PA CTD-binding mutants. Luminescence due to luciferase reconstitution was measured and the signals are represented as a percentage of PA WT. Right Y-axis (logarithmic scale): FluPol PA CTD-binding mutant activity (hatched bars) was assessed by vRNP reconstitution in HEK-293T cells upon transient transfection, using a model vRNA encoding the Firefly luciferase. Luminescence was measured, normalised to a transfection control and is represented as a percentage of PA WT. *p < 0.033, **p < 0.002, ***p < 0.001 (one-way ANOVA; Dunnett’s multiple comparisons test). **(D)** WSN vRNPs with the indicated PA mutations were reconstituted in HEK-293T cells by transient transfection using the NA vRNA segment. Steady-state levels of NA mRNA, cRNA and vRNA were quantified by strand-specific RT-qPCR (Kawakami et al., 2011) and are presented as ratios of mRNA to vRNA (grey bars) or cRNA to vRNA (blue bars) levels relative to PA WT. The corresponding mRNA, vRNA and cRNA levels are presented in Fig. S1E. ***p < 0.001 (one-way ANOVA; Dunnett’s multiple comparisons test). **(E)** FluPol trans-complementation of a transcription-defective (FluPol_(T-)_, PA D108A, (Dias et al., 2009)) and a replication-defective (FluPol_(R-)_, PA K664M, Drncova et al., unpublished) FluPol was measured by vRNP reconstitution in HEK-293T cells upon transient transfection, using a model vRNA encoding the Firefly luciferase. Luminescence was measured at 48 hpt and normalised to a transfection control. The data are represented as percentage of PA WT. **p < 0.002 (one-way repeated measure ANOVA; Dunnett’s multiple comparisons test). **(F-G)** WSN vRNPs with the indicated PA CTD-binding mutations were trans-complemented with (F) a replication-defective (FluPol_(R-)_, PA K664M) or (G) a transcription-defective (FluPol_(T-)_, PA D108A, (Dias et al., 2009)) FluPol. The FluPol_(R-)_ and FluPol_(T-)_-mediated fold-change (FC) relative to the background (x-axis, represented separately in Fig. S1F-G) is plotted against the fold-decrease of viral m-, c- and vRNA levels relative to PA WT as measured in a vRNP reconstitution assay (y-axis, represented separately in Fig. S1E).

To assess this dual role hypothesis, we examined whether binding of the A/WSN/1933 (WSN) FluPol to ANP32A is altered by previously described FluPol CTD-binding mutations (PA K289A, R454A, K635A, or R638A) (Lukarska et al., 2017). For this purpose, we made use of cell-based *G. princeps* split-luciferase protein-protein complementation assays as described before (Mistry et al., 2020; Morel et al., 2022). As expected from our previous work (Lukarska et al., 2017), steady-state levels of the wild-type and mutant PA protein are similar or slightly reduced in the case of the PA K635A mutant (**Fig. S1B**) and each mutation reduces FluPol binding to the CTD (**Fig. 1C**, grey bars) as well as FluPol activity as measured in a vRNP reconstitution assay (**Fig. 1C**, hatched bars). Remarkably, they also reduce FluPol binding to ANP32A (**Fig. 1C**, blue bars). We then tested double mutations (PA K635A+R638A, K289+R454A) or mutations shown to disrupt CTD-binding site 2A more severely (PA S420E, Δ550-loop, highlighted in orange in **Fig S1A**) (Krischuns et al., 2022). Steady-state levels of the mutant PA proteins remain unchanged (**Fig. S1C**) while stronger reductions in ANP32A-binding and FluPol activity are observed (**Fig. S1D,** blue and hatched bars). CTD-binding signals remain in the comparable range of 25-50% signal reduction as with the initial single mutations (**Fig. S1D,** grey bars) indicating that disruption of a single CTD-binding site (site 1A or site 2A) can largely preserve the CTD-FluPol interaction signal through binding to the remaining second FluPol CTD-binding site.

We then quantified the accumulation of viral RNA species (NA mRNA, cRNA, and vRNA) in the vRNP reconstitution assay by strand-specific RT-qPCR (Kawakami et al., 2011). The PA K289A, R454A, K635A, and R638A mutants show a ∼80% and ∼50% decrease of mRNA:vRNA and cRNA:vRNA ratios, respectively, indicating that both transcription and replication are affected (**Fig. 1D**). The absolute levels of cRNA and vRNA are decreased for the PA R454A and K635A mutants, as well as with the PA S420E and Δ550-loop mutants (**Fig. S1E**). Intriguingly, in the presence of the PA K289A and R638A mutants, cRNA and vRNA levels appear unchanged or even increased (**Fig. S1E**). As the vRNP reconstitution assay was shown to be skewed towards transcription (Robb et al., 2009), it is likely that although a replication defect exists, it is masked by the transcription defect as more FluPols become available for replication activity, especially when the defect in ANP32 binding is moderate as observed for the PA R638A mutant (**Fig. 1C**).

To further support the notion that the FluPol CTD-binding interface is essential for FluPol replication, we made use of transcription-defective (FluPol_(T-)_, PA D108A, (Dias et al., 2009) and replication-defective (FluPol_(R-)_, PA K664M, Drncova et al., unpublished) FluPol mutants that trans-complement each other in a vRNP reconstitution assay (**Fig. 1E**). FluPol CTD-binding mutants were either co-expressed with the FluPol_(R-)_ mutant to reveal a defect in transcription (**Fig. S1F**), or with the FluPol_(T)_ mutant to reveal a defect in replication (**Fig. S1G**). Upon co-expression with the FluPol_(R-)_, trans-complementation is observed for most of the investigated CTD-binding mutants. Moreover, the FluPol_(R-)_-mediated fold-change (FC) shows a strong positive correlation when plotted against the reduction in mRNA levels, confirming that the tested FluPol mutants are transcription-defective (**Fig. 1F**). Similarly, upon co-expression with the FluPol_(T-)_, trans-complementation is observed for most of the investigated mutants, with positive correlations being observed between the FluPol_(T-)_-mediated FC and the reduction of cRNA as well as vRNA levels (**Fig. 1G**) demonstrating that the investigated FluPol mutants are defective for replication.

Overall, our findings show that the FluPol CTD-binding interface, beyond its function for FluPol “cap-snatching”, is also involved in the FluPol-ANP32 interaction and is essential for genome replication. They are not limited to the WSN strain, as a dual-effect of the mutations PA K289A, R454A, K635A, or R638A on CTD- and ANP32A-binding is also observed with the FluPol of the human isolate A/Anhui/01/2013 (H7N9) (Anhui-H7N9), a virus of avian origin which harbours the human-adaptation signatures PB2 K627 and N701 (**Fig. S1H**). Sequence alignments of the PB1, PB2, PA and NP proteins from the influenza A viruses used in this study are shown in **Fig. S2**.

### Serial passaging of FluPol PA K289A, R454A, K635A, R638A mutant viruses selects for adaptive mutations which restore FluPol binding to the CTD and ANP32A

We previously rescued by reverse genetic procedures recombinant WSN PA mutant IAVs (PA K289A, R454A, K635A, R638A mutants) that were highly attenuated, formed pinhead-sized plaques and had acquired multiple non-synonymous mutations (Lukarska et al., 2017). In order to investigate whether and how the recombinant viruses adapt to defective CTD and ANP32-binding (**Fig. 1**), we subjected them to eight serial cell culture passages in MDCK-II cells followed by titration and next generation sequencing (NGS) of the viral genome at passage 1 to 4 and 8 (**Fig. 2A**, p1-4 and p8). The plaque size of the passaged viruses progressively increased (**Fig. 2B** and **S3A**), which went along with distinct patterns of adaptive mutations (**Source data file 1**), suggesting that cell culture passaging increased the replicative capacity of the PA mutant viruses. True reversions that require at least two nucleotide substitutions did not emerge. The observed second-site mutations are primarily located on the FluPol subunits as well as the NP (**Fig. 2C**, **S3B** and **S4A**) and are mostly surface exposed (**Fig. 2D** and **S4B-D**). Several FluPol second-site mutations are located close to the FluPol CTD-binding sites (**Fig. 2D**, PA D306N, C489R, R490I, R496Q) while some PB2 second-site mutations are located in flexible hinge-subdomains (**Fig. 2D**, PB2 D253G, S286N, M535I). Overall, the FluPol second-site mutations show no local clustering or common function, indicating that diverse routes of adaptation occurred. NP second-site mutations are also surface exposed, but to our knowledge have not been associated to date with any specific function (**Fig. S4D**) (Tang et al., 2021).

**Figure 2:**
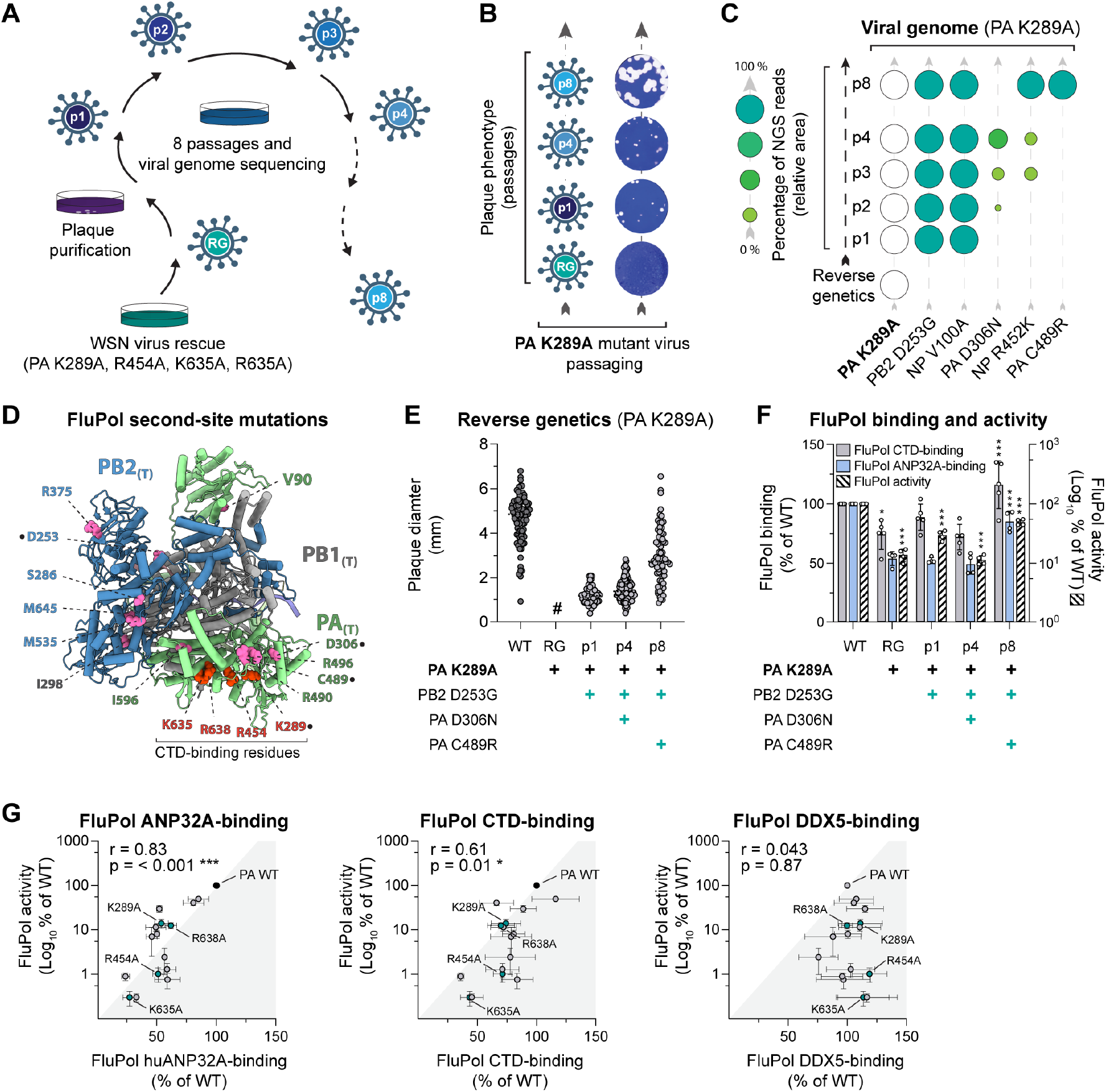
Serial passaging of mutant viruses with mutations at the FluPol-CTD interface selects for adaptive mutations which restore FluPol binding to the CTD and ANP32A. **(A)** Schematic representation of the cell culture passaging experiment. Recombinant WSN mutant viruses (PA K289A, R454A, K635A and R638A) were plaque purified on MDCK-II cells after reverse genetics (p1) and were subjected to seven serial passages in MDCK-II cells (p2 to p8) followed by titration and short-read next generation sequencing (NGS) of the viral genome. **(B)** Plaque phenotype of passaged PA K289A mutant viruses. Representative images of crystal violet stained plaque assays of the supernatants collected after reverse genetics (RG), p1, p4 and p8 are shown. **(C)** The viral genome of passaged PA K289A mutant viruses was sequenced by NGS after p1 - p4 and p8 (from bottom to top). The initial FluPol CTD-binding mutation (PA K289A, white circles) and the second site mutations on PB1, PB2, PA and NP (green circles) are indicated. Only second-site mutations found in ≥ 10% of reads in at least one passage are shown. The fraction of reads showing a given mutation is indicated by the area of the circle, and by different shades of green that represent <25, 25-50, 50-75 and >75 % of reads (schematic on the left). **(D)** Representation of the Cryo-EM structure of the FluPol heterotrimer in the FluPol_(T)_ conformation, bound to a 3’5’ vRNA promoter and a short capped RNA primer (A/NT/60/1968, PDB: 6RR7, (Fan et al., 2019)). Ribbon diagram representation with PA in green, PB1 in grey, and PB2 in blue. Initially mutated FluPol PA residues (PA K289A, R454A, K635A and R638A) are highlighted in red and residues corresponding to FluPol second-site mutations in pink. Second-site mutations which appeared during passaging of the WSN PA K289A mutant virus are highlighted with a dot. **(E)** Characterisation of recombinant WSN mutant viruses. Recombinant viruses with the primary FluPol mutation PA K289A and the indicated combinations of FluPol second-site mutations (PB2 D253G, PA D306N, PA C489R) were generated by reverse genetics. The supernatants of two independently performed RG experiments were titrated on MDCK cells and stained by crystal violet. Plaque diameters (mm) were measured and each dot represents one viral plaque. Representative plaque assays are shown in Fig. S3C. (#) not measurable pinhead-sized plaques. **(F)** The indicated combinations of WSN FluPol second-site mutations which occurred during passaging of the PA K289A mutant virus were characterised in cell-based FluPol binding and activity assays. Left Y-axis (linear scale): FluPol binding to the CTD (grey bars) and ANP32A (blue bars) was assessed using split-luciferase-based complementation assays as in Fig. 1C. Luminescence signals due to luciferase reconstitution are represented as a percentage of FluPol WT. Right Y-axis (logarithmic scale): FluPol activity (hatched bars) was measured by vRNP reconstitution in HEK-293T cells as in Fig. 1C. Normalised luminescence signals are represented as a percentage of FluPol WT. Stars indicate statistical significance compared to the previous passage. *p < 0.033, ***p < 0.001 (one-way ANOVA; Tukey’s multiple comparisons test). **(G)** For each initial FluPol mutant (PA K289A, R454A, K635A, R638A, green dots) and FluPol genotypes observed upon serial passaging (grey dots), FluPol binding to ANP32A, CTD and DDX5 (x-axis in the left, middle and right panel, respectively) were plotted against the FluPol activity as measured in a vRNP reconstitution assay (y-axis). The FluPol binding and FluPol activity data are represented separately in Fig. 2F for PA K289A and in Fig. S3E for PA R454A, K635A, R638A. *p < 0.033, ***p < 0.001 (Pearson Correlation Coefficient (r)).

We first focused on the reversion pathway of PA K289A mutant virus. Recombinant viruses with combinations of FluPol second site mutations observed along the PA K289A reversion pathway at p1, p4 and p8 recapitulate the progressive increase in plaque size observed during serial passaging (**Fig. 2E and S3C**), demonstrating that the FluPol second-site mutations are leading to the observed increase in viral replicative capacity. PB2 and PA proteins with second-site mutations which occurred during passaging of PA K289A mutant virus accumulate at similar levels compared to the WT proteins (**Fig. S3D**). However, the combinations of FluPol second site mutations result in sequential increases and decreases of CTD-binding, ANP32A-binding and/or FluPol activity (**Fig. 2F**). The highest levels of CTD-binding, ANP32A-binding and FluPol activity are observed for the combination of mutations observed at p8 (PA K289A+PB2 D253G+PA C489R), which goes along with the highest replicative capacity observed for the passaged virus at p8 (**Fig. 2B and 2E**).

The reversion pathways of the PA R454A, K635A and R638A mutant viruses were analysed in the same way (**Fig. S3E**). Across all tested combinations of primary FluPol mutations and second-site mutations that occurred during passaging, FluPol activity shows a strong positive correlation with ANP32A-binding (**Fig. 2G**, left panel) as well as a positive correlation with CTD-binding (**Fig. 2G**, middle panel) but not with FluPol-binding to other known cellular partners, such as DDX5 (**Fig. 2G**, right panel) or RED (**Fig. S3F)**.

Overall, we show that serial cell culture passaging of PA K289A, R454A, K635A, and R638A mutant viruses selects for adaptive mutations that rescue viral polymerase activity by restoring FluPol binding to the CTD and/or ANP32A, which is in agreement with our initial observation that the investigated FluPol mutants are impaired for CTD-binding as well as ANP32A-binding (**Fig. 1**).

### *De novo* FluPol replication activity is enhanced *in vitro* in the presence of CTD and ANP32A

To further document a potential interplay between CTD and ANP32-binding to FluPol, we tested the effect of mCherry-CTD (CTD-WT) overexpression on FluPol binding to ANP32A as well as FluPol dimerisation in cultured cells. A FluPol binding-deficient CTD in which all serine 5 residues were replaced with alanines was used as a specificity control (CTD-S5A) (Krischuns et al., 2022). mCherry-CTD-WT and -CTD-S5A accumulate to similar levels (**Fig. S5A**), however only mCherry-CTD-WT leads to a dose-dependent increase of FluPol ANP32A-binding, suggesting that the CTD facilitates FluPol-binding to ANP32 (**Fig. 3A**). CTD overexpression also leads to a dose-dependent increase in WSN FluPol dimerisation (**Fig. 3B**), as measured in a split-luciferase complementation assay (Chen et al., 2019). A similar increase was also observed with the Anhui-H7N9 and B/Memphis/13/2003 FluPols (**Fig. S5B**). Various oligomeric FluPol species that have been described (Carrique et al., 2020; Chang et al., 2015; Fan et al., 2019). There is evidence that the split-luciferase-based dimerisation assay, when FluPol is over-expressed in the absence of any other overexpressed viral or cellular protein, detects mostly the symmetrical FluPol dimer species (Chen et al., 2019; Fan et al., 2019). Here however, we observed that the increase in FluPol dimerisation upon CTD-WT overexpression is less pronounced in cells depleted for ANP32A and ANP32B (ANP32AB KO) (**Fig. S5C-D**), which strongly suggests that i) the split-luciferase-based assay also detects the ANP32-dependent asymetrical dimer species, and ii) CTD overexpression enhances the formation of ANP32AB-dependent dimers. The effect of ANP32A and ANP32B depletion is significant for Anhui-H7N9 and B/Memphis/13/2003 FluPols but not for WSN FluPol (**Fig. S5C**), suggesting that other proteins, e.g. ANP32E, can support CTD-dependent dimer-formation for WSN FluPol (Zhang et al., 2020).

**Figure 3:**
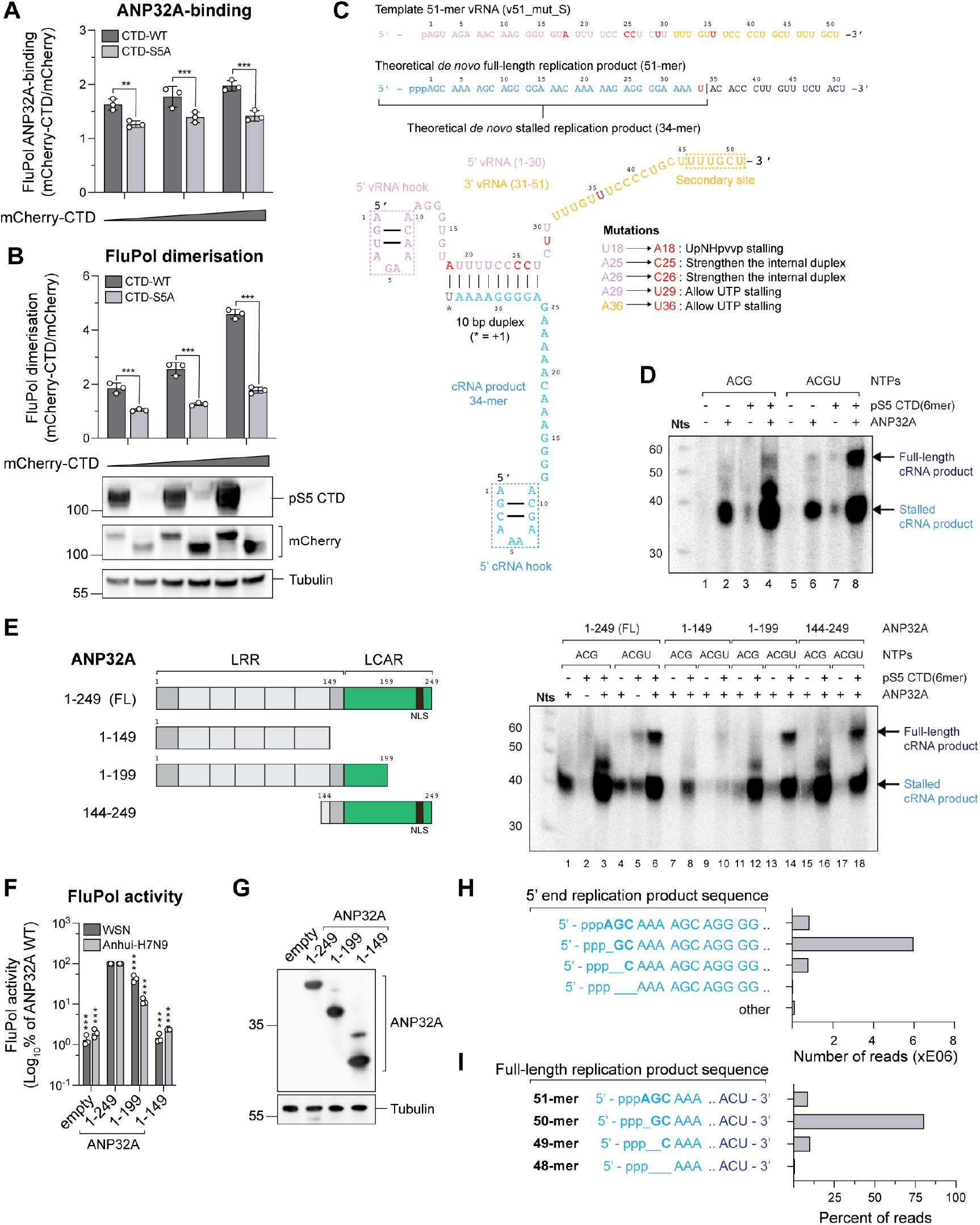
FluPol replication activity is enhanced in the presence of CTD and ANP32A. **(A-B)** Cell-based assays for (A) WSN FluPol-binding to ANP32A and (B) WSN FluPol dimerisation in the presence of increasing amounts of pS5 CTD. HEK-293T cells were co-transfected with expression plasmids for ANP32A tagged with one fragment of the *G. princeps* luciferase and FluPol tagged with the other fragment. FluPol dimerisation was assessed using split-luciferase-based complementation assays as described previously (Chen et al., 2019). Plasmids encoding untagged mCherry or mCherry tagged to CTD-WT or CTD-S5A in which all serine 5 residues were replaced with alanines were co-transfected in increasing amounts. Luminescence due to luciferase reconstitution was measured and the signals are represented as fold-changes compared to untagged mCherry co-expression. In parallel, cell lysates of the FluPol dimerisation assay were analysed by western blot using antibodies specific for the pS5 CTD, mCherry and tubulin. ***p < 0.001 (two-way ANOVA; Sidak’s multiple comparisons test). **(C)** Zhejiang-H7N9 FluPol vRNA loop template design strategy. The 51-mer vRNA template (v51_mut_S) used in *in vitro* replication activity assays and expected theoretical RNA replication products sequences are displayed. The v51_mut_S sequence is derived from segment 4 of A/Zhejiang/DUID-ZJU01/2013(H7N9)/KJ633805. The 5’ vRNA end (1-30) is coloured in pink. The 3’ vRNA end (31-51) is coloured in gold. v51_mut_S introduced mutations are in bold, coloured in red and their functions are summarised. The theoretical *de novo* full-length replication product sequence is coloured in light blue (1-34) and dark blue (35-51). The expected stalled *de novo* replication product sequence using non-hydrolysable UpNHpp is annotated with U34 in red. A schematic representation of the expected stalled elongation state is displayed and coloured accordingly. The 5’ end vRNA hook, the 5’ end cRNA hook of the replication product and the secondary site are highlighted by a dotted rectangle. **(D)** The *de novo* replication activity of Zhejiang-H7N9 FluPol using v51_mut_S is enhanced in the presence of both ANP32A and pS5 CTD. The assays are done in the presence of either 3 NTPs (AUG) or 4 NTPs (AUGC), with or without pS5 CTD and ANP32A. Full-length and stalled replication products are indicated on the right side of the gel and coloured as in C. The decade molecular weight marker (Nts) is shown on the left side of the gel. Uncropped gels are provided as a source data file. **(E)** (left) Schematic representation of the full-length (FL, 1-249) ANP32A and the investigated deletion mutants: ANP32A 1-149, 1-199, 144-249. LRR: leucine-rich repeat; LCAR: low-complexity acidic region; NLS: nuclear localization signal. (right) The *de novo* replication activity of Zhejiang-H7N9 FluPol using v51_mut_S is dependent on both the LRR and LCAR domains of ANP32A. The assays are done in the presence of either 3 NTPs (AUG) or 4 NTPs (AUGC), with or without pS5 CTD and with either FL ANP32A or the deletion mutants represented in (E). *De novo* replication products are annotated on the right side of the gel and coloured as in C. The decade molecular weight marker (Nts) is shown on the left side of the gel. Uncropped gels are provided as a source data file. **(F)** WSN (dark grey bars) and Anhui-H7N9 (light grey bars) vRNPs were reconstituted in HEK-293T ANP32AB KO cells with a model vRNA encoding the Firefly luciferase, and co-expressed with either the FL ANP32 or the indicated deletion mutants, tagged at their C-terminus to a V5 tag and an SV40 nuclear localisation sequence. Luminescence was measured and normalised to a transfection control. The data are represented as a percentage of ANP32A FL. ***p < 0.001 (two-way ANOVA; Sidak’s multiple comparisons test). **(G)** HEK-293T cells were transfected with the indicated ANP32A expression plasmids. Cells lysates were analysed by western blot using antibodies specific for the V5-tag and tubulin. Uncropped gels are provided as a source data file. **(H)** RNA-sequencing analysis of Zhejiang-H7N9-4M FluPol *de novo* replication products in the presence of all NTPs, the pS5 CTD and ANP32A. The number of reads is plotted according to the recurrence of the exact 5’ cRNA replication product motifs indicated on the left. Other reads that do not encompass these motifs are plotted as “Other”. NGS data are provided as a source data file. **(I)** RNA-sequencing analysis of FluPol Zhejiang-H7N9-4M *de novo* replication products in the presence of pS5 CTD and ANP32A. The percentage of full-length replication products are plotted according to the 5’ terminal nucleotide which is indicated on the left. NGS data are provided as a source data file.

To investigate by biochemical and structural approaches whether and how *in vitro* replication is dependent on ANP32A and pS5 CTD, we performed unprimed RNA synthesis assays using recombinant A/Zhejiang/DTID-ZJU01/2013(H7N9) FluPol (Kouba et al., 2023) bound to a 51-mer vRNA loop template (denoted v51_mut_S). This polymerase, referred to below as Zhejiang-H7N9, originates from a human isolate of an avian strain possessing the characteristic PB2 E627 and N701 residues (**Fig. S2**) (Chen et al., 2013). The sequence of the v51_mut_S template was derived by joining of the first 30 and the last 21 nucleotides of the respective 5’ and 3’ vRNA ends of Zhejiang-H7N9 segment 4, and was adapted so that a stalled elongation product of 34 nts could be produced by replacing UTP in the reaction by non-hydrolysable UpNHpp (**Fig. 3C**). This would produce a 34-mer cRNA product, estimated to be long enough that its 5’ hook (1-10) could bind to the encapsidase according to the modelled Zhejiang-H7N9 replicase-encapsidase structure. The addition of all NTPs would produce a full-length 51-mer replication product without poly-adenylation (A17, only 4xU), assuming RNA synthesis proceeded to the 5’ end of the template, rather than terminating prior to reading through the 5’ hook.

*De novo* replication assays were performed with WT Zhejiang-H7N9 FluPol, using three (ACG) or four (ACGU) nucleotides, (a) excess apo-polymerase, that could serve as encapsidase, (b) huANP32A and (c) 6-repeats pS5CTD peptide (**Fig. 3D**, pS5 CTD(6mer)). The results show that in reactions with ACG only (**Fig. 3D**, lane 1-4), the principal product is the stalled 34-mer cRNA, whose significant production is only visible in the presence of CTD and ANP32A (**Fig. 3D**, lane 4). When all four nucleotides are added (**Fig. 3D**, lane 5-8), the presence of CTD and ANP32A again strongly enhances RNA synthesis with the production of both stalled product (34-mer) and a significant amount of full-length replication product (**Fig. 3D**, lane 8).

We further investigated which domains of huANP32A are essential for this *de novo* replication activity increase in the presence, or not, of pS5 CTD(6mer) peptide. By comparing the full-length ANP32 (“FL” lane 1-6) with ANP32A subdomains corresponding to the LRR (“1-149” lane 7-10), the LRR and LCAR N-terminal region (“1-199” lane 11-14), or the LCAR and LRR C-terminal region (“144-249” lane 15-18) (**Fig. 3E**, left panel), we show that the 149-199 region of ANP32A is most critical to observe a joint CTD- and ANP32-dependent enhancement of replication product synthesis (**Fig. 3E**, right panel). These *in vitro* data are consistent with cell-based assays in which FluPol activity was investigated with the same C-terminal deletion mutants of ANP32A (**Fig. 3F and 3G**).

To confirm whether a 5’ triphosphorylated full-length replication product was actually synthesised, we performed next-generation sequencing (NGS) of all RNAs present in the reaction mix (**Fig. 3D**, lane 8). Prior to NGS and to simplify the analysis of this heterogeneous RNA sample, we first degraded non-5’ triphosphorylated RNAs with a 5’-phosphate-dependent exonuclease. This removes the v51_mut_S template, which carries a 5’ monophosphate, and which is perfectly complementary to the expected product and could have formed unwanted double-stranded RNA (**Fig. S5E**). NGS analysis revealed that the majority of reads (*i.e.*, premature and near full-length products) start with 5’-ppp**<u>_G</u>**C… instead of the expected 5’-ppp**<u>A</u>**GC… (**Fig. 3H**). Despite the presence of many premature termination products (**Fig. S5F**), around 80% of near full-length products, are 50-mers, lacking the expected 5’ pppA, with few 51-mers (5’-ppp**<u>A</u>**GC…) and 49-mers (5’-ppp **<u>C</u>**…) (**Fig. 3I**). This suggests that, *in vitro,* most RNA synthesis products are initiated *de novo* with a 5’-pppG at the second position of the v51_mut_S template (3’-U**<u>C</u>**G…).

### Structural analysis shows that both replication and transcription are consistent with CTD-binding to the FluPol

Using cryo-EM, we then sought to determine structures of actively replicating FluPol, in the presence of ANP32A, pS5 CTD(6mer) peptide mimic and excess apo-FluPol, by freezing grids with the *de novo* reaction stalled with UpNHpp (**Fig. 3D**, lane 4). However, instead of using the WT Zhejiang-H7N9 FluPol, which forms a robust symmetrical dimer *in vitro* (Krischuns et al., 2022) that could compete with the formation of an active asymmetric FluPol_(R)_-FluPol_(E)_ dimer, we used a Zhejiang-H7N9 FluPol mutant that harbours four mutations (PA E349K, PA R490I, PB1 K577G, and PB2 G74R) and is referred to below as the 4M mutant. PA E349K, PB1 K577G, and PB2 G74R were shown to disrupt the symmetrical apo FluPol dimer interface (Chen et al., 2019), while PA R490I appeared during passaging of the PA R638A mutant virus and increases FluPol-ANP32A binding in a cell-based assay (**Fig. S3E**). Indeed, Zhejiang-H7N9 4M shows enriched monomeric particles while preserving a similar *in vitro* activity when compared to the Zhejiang-H7N9 WT FluPol (**Fig. S6A-B)**. Accordingly, in cell-based assays Anhui-H7N9 4M shows decreased FluPol dimerisation, increased binding to ANP32A, while showing activity levels comparable to the WT Anhui-H7N9 FluPol as measured in a vRNP reconstitution assay (**Fig. S6C**).

Several high-resolution structures were obtained from a Krios data collection (**Table 1a**, **Fig. 4, Fig. S7**) including FluPol with PB2-C in a replicase-like configuration at 2.9 Å resolution (**Fig. 4A**, left panel). However, the PA endonuclease domain (PA-ENDO) is still in an unrotated transcriptase-like conformation, thus precluding the usual interaction with the PB2-NLS domain, as previously observed for H5N1 or Flu_C_Pol replicase conformations (**Fig. 4A**, middle and right panel) (Carrique et al., 2020; Fan et al., 2019). A second structure, obtained at a higher resolution (2.5 Å), is in a similar state, but PB2-C is not visible (**Fig. 4B**). For both structures, FluPol is in the pre-initiation state mode A, with the 3’ end of the v51_mut_S template (3’-UCG…) in the active site and the priming loop fully extended and well ordered (**Fig. 4C, D; Fig. S8A)**. The template is positioned with C2 and G3 respectively at the −1 and +1 positions, whilst only the phosphate of U1 is visible (**Fig. 4C-D)**. Finally, a stalled elongation state was obtained at 2.9 Å resolution in which FluPol encloses a template-product duplex, with incoming UpNHpp at the +1 active site position, and the 3′ vRNA end bound to the secondary site (**Fig. 4E, Fig. S8B**). In all structures, the pS5 CTD is observed bound in sites 1A and 2A. Continuous density between the two sites, with 25 pS5 CTD residues coming from four repeats, is observed for the pre-initiation state and the elongation structures (**Fig. 4F**), whereas for the replicase-like structure, the connectivity is lost between both sites. Despite extensive cryo-EM data analysis, ANP32A was never visualised nor the putative replicase-encapsidase dimer.

**Table 1a.** Cryo-EM structures data collection, refinement and validation statistics.

| Name of structure | Influenza A/H7N9 polymerase in replicase-like conformation in pre-initiation state with Pol II pS5 CTD peptide mimic bound in site 1A/2A | Influenza A/H7N9 polymerase in pre-initiation state with continuous Pol II pS5 CTD peptide mimic bound in site 1A/2A | Influenza A/H7N9 polymerase in elongation state with continuous Pol II pS5 CTD peptide mimic bound in site 1A/2A |
| --- | --- | --- | --- |
| PDB ID | PDB ID 8PM0 | PDB ID 8PNP | PDB ID 8PNQ |
| EMDB ID | EMD-17755 | EMD-17782 | EMD-17783 |
| **Data collection and processing** |  |  |  |
| Microscope | ThermoFisher Krios TEM |  |  |
| Voltage (kV) | 300 |  |  |
| Camera | Gatan K3 direct electron detector mounted on a Gatan Bioquantum LS/967 energy filter |  |  |
| Magnification | 105000 |  |  |
| Nominal defocus range ($\mu\text{m}$) | -0.8 / -2.0 | | |
| Electron exposure ($\text{e}^-/\text{\AA}^2$) | 40 | | |
| Number of frames collected (no.) | 40 |  |  |
| Number of frames processed (no.) | 40 |  |  |
| Pixel size ($\text{\AA}$) | 0.84 | | |
| Initial micrographs (no.) | 6000 |  |  |
| Final micrographs (no.) | 5883 |  |  |
| **Refinement** |  |  |  |
| Particles per class (no.) | 33395 | 260565 | 14878 |
| Map resolution ($\text{\AA}$), 0.143 FSC | 2.9 | 2.5 | 2.9 |
| Model resolution ($\text{\AA}$), 0.5 FSC | 3.0 | 2.5 | 2.9 |
| Map sharpening $B$ factor ($\text{\AA}^2$) | -20 | -30 | -20 |
| Map versus model cross-correlation (CCmask) | 0.8428 | 0.8901 | 0.8826 |
| **Model composition** |  |  |  |
| Non-hydrogen atoms | 16673 | 14464 | 14917 |
| Protein residues | 1997 | 1716 | 1727 |
| Nucleotide residues | 31 | 29 | 45 |
| Water | - | 43 | - |
| Ligands | 1 Mg | - | 2 Mg, 1 UpNHpp |
| **B factors (Å<sup>2</sup>)** |  |  |  |
| Protein | 74.00 | 67.24 | 70.02 |
| Nucleotide | 72.82 | 60.96 | 83.39 |
| Ligand | 48.38 | - | 68.99 |
| Water | - | 33.97 | - |
| **R.M.S. deviations** |  |  |  |
| Bond lengths (Å) | 0.003 | 0.003 | 0.005 |
| Bond angles (°) | 0.475 | 0.499 | 0.465 |
| **Validation** |  |  |  |
| MolProbity score | 1.53 | 1.35 | 1.40 |
| All-atom clash score | 4.79 | 4.32 | 3.86 |
| Poor rotamers (%) | 2.03 | 1.24 | 2.01 |
| **Ramachandran plot** |  |  |  |
| Favored (%) | 97.77 | 97.70 | 98.24 |
| Allowed (%) | 2.23 | 2.24 | 1.76 |
| Outliers (%) | 0.00 | 0.06 | 0.00 |

**Figure 4:**
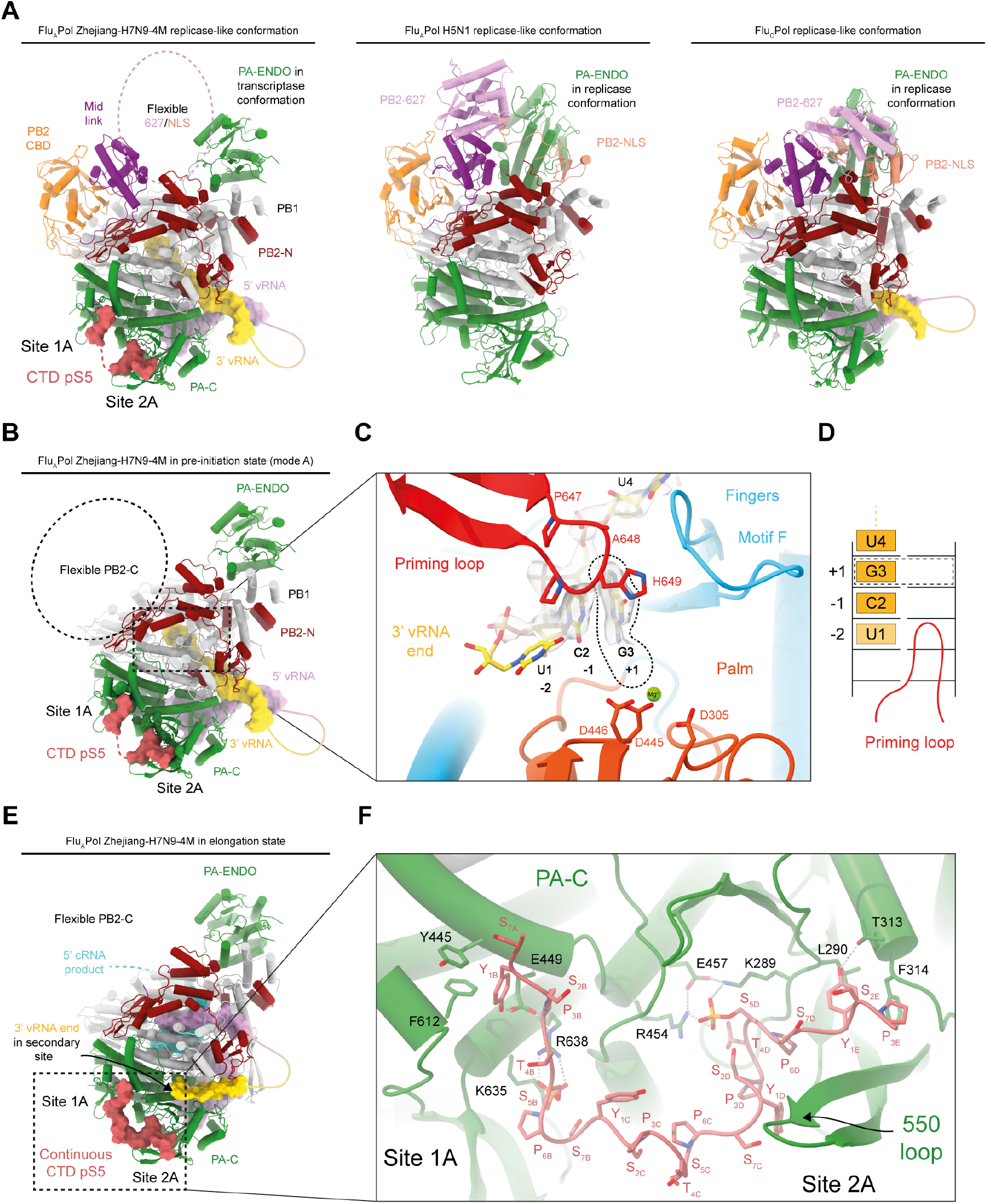
FluPol replication is consistent with CTD-binding to the FluPol. **(A)** Cartoon representations of Flu_A_Pol Zhejiang-H7N9-4M in a replicase-like conformation bound to CTD mimicking-peptides (obtained in this study) and as a reference, previously published Flu_A_Pol replicase-like (A/duck/Fujian/01/2002(H5N1), PDB: 6QPF (Fan et al., 2019)) and the Flu_C_Pol replicase-like conformation extracted from the Flu_C_Pol_R_-Flu_C_Pol_E_-ANP32A asymmetric dimer (C/Johannesburg/1/1966, PDB: 6XZG, (Carrique et al., 2020)). FluPol subunits and domains are coloured as follow: PA subunit in green, PB1 subunit in light grey, PB2-N in dark red, PB2 cap-binding domain (PB2-CBD) in orange, PB2 mid-link in pink, PB2 627 in light violet and PB2 NLS in salmon. The pS5 CTD bound to FluPol Zhejiang-H7N9-4M in site 1A and 2A is coloured in red and displayed as surface. The pS5 CTD discontinuity between both sites is shown as a dotted line. The flexibility of Flu_A_Pol Zhejiang-H7N9-4M PB2-627/NLS domains is highlighted as a dotted circle. The three displayed FluPols are in the same orientation, aligned on the PB1 subunit. In the Flu_A_Pol Zhejiang-H7N9-4M structure the PA-Endonuclease domain (PA-ENDO) remains in a transcriptase conformation, while it adopts a replicase conformation, interacting with the PB2-NLS domain, in the FluPol Fujian-H5N1 and Flu_C_Pol structures. RNAs are displayed as surfaces. The 5’ vRNA end is coloured in pink and the 3’ vRNA end in yellow. Flexible nucleotides are represented as solid line. **(B)** Cartoon representation of FluPol Zhejiang-H7N9-4M structure in the pre-initiation state (mode A), obtained at 2.5 Å resolution by cryo-EM. The colour code is the same as in A. PB2 C-terminal domains (PB2-C) are flexible and highlighted as a dotted circle. **(C)** Close-up view on the 3’ vRNA end template (v51_mut_S) position in FluPol Zhejiang-H7N9-4M active site. The Coulomb potential map of the template is shown. The 3’ vRNA end is coloured in yellow, by heteroatom. The 3’ vRNA terminal U1 is displayed but remains unseen in the cryo-EM map. The 3’ vRNA G3 is in the +1 active site position, highlighted by a dotted line. The priming loop is coloured in red and tip residues are shown (P647, A648, H649, P650). The palm domain is coloured in orange and the catalytic aspartic acids (D445, D446 and D305) are displayed, coordinating a Mg^2+^ ion, coloured in green. The finger domain is coloured in blue and the beta-hairpin motif F is annotated. **(D)** Schematic representation of the 3’ vRNA end terminal nucleotides according to their active site position, as seen in C. 3’-U1 is in position “-2”, C2 in position “-1”, G3 in position “+1”. The +1 active site position is highlighted by a dotted rectangle. The priming loop position is indicated. **(E)** Cartoon representation of FluPol Zhejiang-H7N9-4M in a stalled elongation state. The colour code is the same as in A. PB2-C domains are flexible. The visible part of the cRNA *de novo* replication product is displayed as surface, while the flexible part is shown as a dotted line. The 3’ vRNA end is bound to the secondary site. The CTD pS5 is continuous between sites 1A and 2A. **(F)** Close-up view on the pS5 CTD interaction with FluPol Zhejiang-H7N9-4M PA-C domain. FluPol residues interacting with the pS5 CTD are shown for sites 1A and 2A. Each CTD pS5 repeat is indicated (S_7A_ | Y_1B_S_2B_P_3B_T_4B_S_5B_P_6B_S_7B_ | Y_1C_S_2C_P_3C_T_4C_S_5C_P_6C_S_7C_ | Y_1D_S_2D_P_3D_T_4D_S_5D_P_6D_S_7D_ | Y_1E_S_2E_P_3E_). The PA-550 loop is annotated.

Altogether, these results show that *in vitro*, *de novo* RNA synthesis assays result in a heterogeneous mix of FluPol conformations including replicase-like and transcriptase-like configurations, reflecting the flexibility of the peripheral PB2-C domains (**Fig. S7**). Importantly, a replicase-like initiation complex is observed for the first time, characterised by the radically different position of the cap-binding domain compared to the transcriptase conformation (**Fig. 4A**) (Thierry et al., 2016), which has never been observed during extensive studies of cap-dependent transcription (Wandzik et al., 2020). However, the observed stalled elongation state, in which the translocated 3’ end of the template has reached the secondary binding site (**Fig. 4E**), is similar to the previously described pre-termination transcription state (Wandzik et al., 2020). Finally, all structures have the CTD peptide mimic bound in both sites 1A and 2A, suggesting that both replication and transcription are consistent with CTD binding (**Fig. 4A-B, E**).

The pre-initiation state structures show that at least *in vitro*, and using highly purified recombinant FluPol, the preferred position of the 3’ end of the vRNA template is with C2, rather than U1, at the −1 position. This leads to formation of pppGpC at the beginning of the product, consistent with the NGS results, or alternatively, would allow efficient priming by pppApG, if this dinucleotide was already available from another source. Internal initiation at position 2 of vRNA has been previously described (Vreede and Brownlee, 2007; Zhang et al., 2010), but other authors report pppApG formation at position 1 (Deng et al., 2006). We note that initiation at position 2 of vRNA implies that the 5’ end of the cRNA product probably forms a less stable hook structure since the A1:A10 non-canonical base pair would be disrupted. This could explain why we do not observe in cryo-EM the cRNA hook bound in a putative encapsidase, even though there is excess apo-polymerase. This could further explain why the putative asymmetric FluPol_(R)_-FluPol_(E)_ dimer is also not observed, since it seems likely that *in vitro*, the Flu_A_Pol FluPol_(R)_-FluPol_(E)_ -ANP32 complex is not stable in itself, unlike the case of Flu_C_Pol, where the FluPol_(R)_-FluPol_(E)_ complex is stable even in the absence of ANP32 (Carrique et al., 2020). These observations suggest that other cellular or viral factors and the RNP context might be required to recapitulate true terminal initiation of vRNA replication and further work is required to clarify this issue (Drncova et al., unpublished).

### Restoration of CTD-binding function enhances FluPol binding to ANP32A and FluPol replication activity

Second-site mutations PA C489R and PB2 D253G were selected during passaging of the recombinant WSN virus with a PA K289A mutation (**Fig. 2C**) and conferred elevated FluPol-binding to the CTD as well as to ANP32A (**Fig. 2F**, grey and blue bars, respectively). PA C489 is in close proximity to the phosphoserine binding site in CTD-binding site 2A (**Fig. 5A**). The mutation PA C489R could therefore plausibly compensate for the loss of the positive charge of PA K289A and rescue the interaction with the CTD. Indeed, structural analysis by cryo-EM of Zhejiang-H7N9 WT FluPol in the symmetric dimeric form (**Table 1b** and **Fig. S9**) and bearing the double mutation K289A+C489R confirms that C489R points towards the phosphoserine binding site, although at a slightly greater distance than K289 (**Fig. 5B-C).** Despite this, the structure did not reveal CTD binding to the Zhejiang-H7N9 PA K289A+C489R FluPol in this conformation. Therefore, we sought to analyse the functional impact of the single PA C489R reversion on the viral phenotype, and in cellular assays, on FluPol activity and FluPol-binding to the CTD and ANP32A (**Fig. 5D-I**).

**Figure 5:**
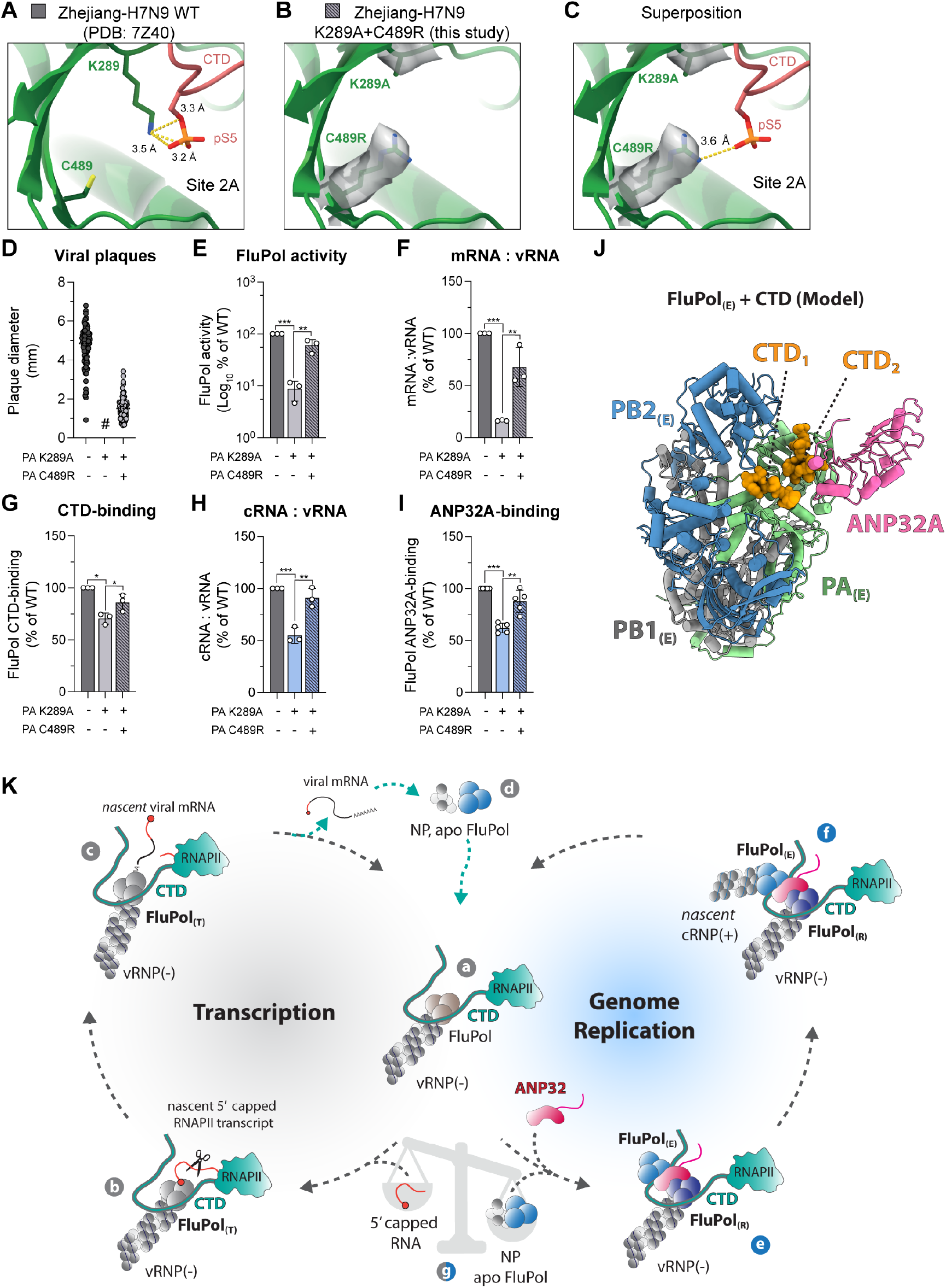
Restoration of CTD-binding enhances FluPol binding to ANP32A and FluPol replication activity - a model for RNAP II CTD-anchored transcription and replication of the influenza virus genome. **(A)** Cartoon representation of the pS5 CTD bound to FluPol Zhejiang-H7N9 in site 2A (PDB: 7Z4O, (Krischuns et al., 2022)). The PA subunit is coloured in green and the pS5 CTD is coloured in red. PA K289 and PA C489 residues are displayed. Putative hydrogen bonds between PA K289 and pS5 are drawn as yellow dashed lines and the corresponding distances are indicated. **(B)** Cartoon representation of FluPol Zhejiang-H7N9 PA K289A+C489R (obtained in this study). The PA subunit is coloured in green. The Coulomb potential map of PA K289A and C489R is shown and the corresponding residues are displayed. The pS5 CTD is not present. **(C)** Superposition of the pS5 CTD, extracted from the structure shown in A, with the FluPol Zhejiang-H7N9 PA K289A+C489R structure shown in B. The putative hydrogen bond between PA C489R and pS5 is shown and the corresponding distance is indicated. **(D-I)** Phenotypes associated with the WSN FluPol PA K289A mutation and the second-site mutation PA C489R were characterised. (D) Plaque phenotype of recombinant WSN mutant viruses produced by reverse genetics. The supernatants of two independently performed RG experiments were titrated on MDCK cells and stained by crystal violet. Plaque diameters (mm) were measured and each dot represents one viral plaque. Representative plaque assays are shown in Fig. S10A (#) not measurable pinhead-sized plaques. (E) WSN FluPol activity was measured by vRNP reconstitution in HEK-293T cells, using a model vRNA encoding the Firefly luciferase. Luminescence was measured and normalised to a transfection control. The data are represented as a percentage of PA WT. (F) WSN vRNPs were reconstituted in HEK-293T cells by transient transfection using the NA vRNA segment. The steady-state levels of NA mRNA and vRNA were quantified by strand-specific RT-qPCR (Kawakami et al., 2011), normalised to GAPDH by the 2^−ΔΔCT^ method (Livak and Schmittgen, 2001) and are presented as ratios of mRNA to vRNA levels relative to PA WT. The corresponding RNA levels are presented in Fig. S10B. (G) WSN FluPol binding to the CTD was assessed using a split-luciferase-based complementation assay. HEK-293T cells were co-transfected with expression plasmids for the CTD tagged with one fragment of the *G. princeps* luciferase and FluPol tagged with the other fragment. Luminescence due to luciferase reconstitution was measured and the data are represented as a percentage of PA WT. (H) Accumulation levels of cRNA and vRNA in a vRNP reconstitution assay were determined by strand-specific RT-qPCR (Kawakami et al., 2011) as in F. The corresponding RNA levels are presented in Fig. S10B. (I) ANP32A was tagged with one fragment of the *G. princeps* luciferase and binding to WSN FluPol was determined as in G. *p < 0.033, **p < 0.002, ***p < 0.001 (one-way ANOVA; Dunnett’s multiple comparisons test). **(J)** Model of the encapsidating Zhejiang-H7N9 FluPol_(E)_-ANP32 interface. The model was constructed by superposing equivalent Zhejiang-H7N9 Flu_A_Pol domains of 5’ hook bound apo-dimer Zhejiang-H7N9 Flu_A_Pol (PDB: 7ZPL, (Kouba et al., 2023)) for the FluPol_(E)_ on those of Flu_C_Pol replication complex (PDB: 6XZQ, (Carrique et al., 2020)). The ANP32A was left unchanged. The pS5 CTD peptide mimic was added to the model by superposing the H17N10 Flu_A_Pol structure with bound pS5 CTD peptide (PDB:5M3H, (Lukarska et al., 2017)) on the FluPol_(E)_. **(K)** A model for RNAP II CTD-anchored transcription and replication of the influenza virus genome. Upon influenza virus infection, incoming vRNPs are imported into the nucleus and bind to the host RNA polymerase II (RNAP II) C-terminal domain (CTD) through bipartite interaction sites on the influenza virus polymerase (FluPol) (**a**). This intimate association allows FluPol, in the transcriptase conformation (FluPol_(T)_), to cleave short capped oligomers derived from nascent RNAP II transcripts in a process referred to as “cap-snatching” to initiate primary transcription of viral mRNAs (**b**). Polyadenylation is achieved by a non-canonical mechanism involving stuttering of the viral polymerase at a polyadenylation signal (**c**). The 5′ and 3′ vRNA extremities always remain bound to the polymerase which allows efficient recycling from the termination to the initiation state (**c-a**). Upon translation of viral mRNAs, de novo synthesised FluPols in an apo state (not viral RNA-bound) and NPs are imported into the nucleus (**d**). The apo FluPol, in conjunction with the host factor ANP32, associates with the parental CTD-associated FluPol_(T)_ and triggers its conformational transition into a replicating FluPol_(R)_, to form an asymmetric FluPol_(R)_-FluPol_(E)_ dimer (**e**) where FluPol_(E)_ is encapsidating the newly synthesized cRNA in conjunction with NP (**f**). The FluPol_(R)_-FluPol_(E)_-ANP32 replication complex remains associated to the RNAP II through direct binding of the CTD to FluPol_(R)_, FluPol_(E)_ as well as ANP32 throughout synthesis of the progeny cRNP. Anchoring of the parental vRNP to the CTD allows it to engage into successive cycles of either viral genome replication or mRNA transcription, depending on the availability of NP, apo FluPol and/or nascent capped oligomers derived from actively transcribing RNAP II (**g**). Such switching between both activities allows efficient adaption to waving levels of de novo synthesised vRNP components in the nucleus of an infected cell and is key to ensure a correct balance between genome replication and mRNA transcription.

**Table 1b.** Cryo-EM structures data collection, refinement and validation statistics. K289A/C489R revertant.

| **Name of structure** | **A/H7N9 K289A/C489R**<br>**Apo-dimer with promoter** |
| --- | --- |
| PDB ID | PDB ID 8POH |
| EMDB ID | EMD-17792 |
| **Data collection and processing** |  |
| Microscope | ThermoFisher Glacios |
| Voltage (kV) | 200 |
| Camera | Falcon IV/Selectris X |
| Magnification | 130k |
| Nominal defocus range (µm) | -0.8 / -2.0 |
| Electron exposure (e<sup>-</sup>/Å<sup>2</sup>) | 40 |
| Number of fractions processed (no.) | 24 |
| Pixel size (Å) | 0.907 |
| Micrographs (no.) | 5092 |
| **Refinement** |  |
| Particles per class (no.) | 278761 |
| Map resolution (Å), 0.143 FSC | 3.26 |
| Model resolution (Å), 0.5 FSC | 3.38 |
| Map sharpening *B* factor (Å<sup>2</sup>) | -140 |
| Map versus model cross-correlation (CCmask) | 0.7456 |
| **Model composition** |  |
| Non-hydrogen atoms | 20729 |
| Protein residues | 2479 |
| Nucleotide residues | 38 |
| Water | - |
| Ligands | 1 Mg |
| ***B* factors (Å<sup>2</sup>)** |  |
| Protein | 105.84 |
| Nucleotide | 123.86 |
| Ligand | 33.48 |
| Water | - |
| **R.M.S. deviations** |  |
| Bond lengths (Å) | 0.003 |
| Bond angles (°) | 0.562 |
| **Validation** |  |
| MolProbity score | 1.84 |
| All-atom clash score | 6.79 |
| Poor rotamers (%) | 2.37 |
| **Ramachandran plot** |  |
| Favored (%) | 96.91 |
| Allowed (%) | 3.09 |
| Outliers (%) | 0.0 |

A recombinant WSN virus with the PA K289A+C489R mutations shows larger plaques compared to the PA K289A mutant virus, demonstrating that the PA C489R second-site mutation provides a significant growth advantage to PA K289A mutant virus (**Fig. 5D** and **Fig. S10A**). Indeed, WSN FluPol activity as measured in a vRNP reconstitution assay increases significantly when the PA C489R mutation is combined with PA K289A (**Fig. 5E**). This increase correlates with a restored mRNA:vRNA ratio (**Fig. 5F** and **Fig. S10B**) as well as an increased FluPol-binding to the CTD (**Fig. 5G**). Importantly, steady-state levels of the wild-type and mutant PA proteins are similar (**Fig. S10C**). These observations suggest that charge restoration in CTD-binding site 2A of the WSN FluPol in the transcriptase conformation rescues the interaction with the host RNAP II for “cap-snatching” in cell-based assays and during live-virus infection. In cell-based assays, the Anhui-H7N9 FluPols with PA K289A and PA K289A+C489R show similar trends to those observed with their WSN FluPol counterparts, although with much less pronounced phenotypes (**Fig. S10D**). Therefore we speculate that the fact that CTD-binding is not restored *in vitro* for Zhejiang-H7N9 PA K289A+C489R could be due to strain-specificities (**Fig. S2**) and/or to the presence of cellular factors that favor binding of the CTD to the PA K289A+C489R and are missing in the *in vitro* assay.

Beyond the rescue of FluPol transcription, cRNA accumulation levels increase significantly when PA C489R is combined with PA K289A (**Fig. S10B**), which leads to a rescue of the imbalanced cRNA:vRNA ratio associated with the PA K289A mutant (**Fig. 5H**). This goes along with increased ANP32A-binding (**Fig. 5J**). Taken together our observations suggest that the second-site mutation PA C489R has another functional impact, namely restoring FluPol replication by enhancing ANP32A-binding. Similar results are obtained for the PA C453R revertant which has been shown to restore FluPol-binding to the CTD in site 1A by compensating the loss of positive charge of the PA R638A mutation (**Fig. S10E)** (Fodor et al., 2003; Lukarska et al., 2017). PA C453R rescues the attenuated viral plaque phenotype (**Fig. S10F-G**) as well as the reduced FluPol activity (**Fig. S10H**) associated with PA R638A, while it does not affect PA steady-state levels (**Fig. S10J**). Strikingly, PA C453R enhances ANP32A-binding when tested in combination with PA R638A (**Fig. S10I**), again mimicking at CTD-binding site 1A the effects of PA C489R at site 2A.

The positive impact of the PA C489R and PA C453R mutations on ANP32A-binding could possibly be mediated by an increased binding of the pS5 CTD in site 1A or 2A in either the FluPol_(R)_ and/or FluPol_(E)_ conformation. According to our model of the Flu_A_Pol replicase-encapsidase complex (**Fig. 1B**) the FluPol CTD-binding residues on the encapsidase do not mediate a direct interaction between the FluPol_(E)_ and ANP32A. Within the space lined by ANP32A and FluPol_(E)_, it is possible to model CTD repeats, which make contacts with the FluPol_(E)_ CTD-binding sites (1A and 2A) on one side and ANP32A on the other side (**Fig. 5J**). Moreover, Flu_C_Pol X-ray electron density maps revealed an asymmetric FluPol dimer crystal packing which share a similar inter-polymerase interface compared to the ANP32A-bound Flu_C_Pol replication complex, with the exception of PB2 peripheral domains (**Fig. S10K-L**). The observed CTD bound to Flu_C_Pol_(R2)_ in site 2C is mostly compatible with ANP32A binding and in a similar position as in our CTD-bound Flu_A_Pol replication complex model. Therefore, although we cannot rule out that the CTD-binding sites 1A and 2A are involved in direct protein-protein interaction between the FluPol_(E)_ and ANP32A, it seems more likely that CTD repeats are bound in between the FluPol_(E)_ and ANP32 to stabilise the FluPol replication complex thereby facilitating FluPol replication.

## Discussion

### A model for RNAP II CTD-anchored transcription and replication of the influenza virus genome

There is long-standing and extensive evidence that binding of FluPol to the pS5 CTD repeats of host RNAP II is essential for transcription of viral mRNAs (Chan et al., 2006; Engelhardt et al., 2005; Newcomb et al., 2009). Here, we challenge the prevailing concept that CTD binding exclusively stabilises the transcriptase conformation of the influenza virus polymerase (FluPol_(T)_) (Zhu et al., 2023). We show that CTD-binding also enhances replication of the viral genome, in conjunction with the host protein ANP32A, which was recently described to bridge two FluPol moieties in an asymmetric replicase-encapsidase dimer (FluPol_(R)_-FluPol_(E)_) (Carrique et al., 2020). Our findings open new perspectives on the spatial coupling of viral transcription and replication and the coordinated balance between these two activities.

Early in infection, incoming vRNPs serve as a template for both primary mRNA transcription and the first round of cRNA synthesis (Vreede et al., 2004). Based on our findings, we propose a model in which incoming vRNPs switch from primary transcription to cRNA synthesis while remaining bound to the RNAP II CTD (**Fig. 5K, a**). It has been long understood that primary transcription requires FluPol_(T)_ association with the RNAP II CTD to enable “cap-snatching” for the priming of viral mRNA synthesis in the FluPol_(T)_ conformation (**Fig. 5K, b-c**). Our structural and biochemical data reveal that a vRNA- and CTD-bound FluPol can adopt a FluPol_(R)_ conformation and initiate vRNA to cRNA synthesis *de novo*, and that CTD binding to the FluPol enhances cRNA synthesis. Importantly, the switch from a transcribing FluPol_(T)_ to a replicating FluPol_(R)_ can occur only after *de novo* synthesised FluPol and NP are imported into the nucleus (**Fig. 5K, d**) and an asymetric FluPol_(R)_-ANP32A-FluPol_(E)_ complex is assembled, ensuring encapsidation of the nascent viral RNA by the FluPol_(E)_ moiety. Our cell-based and *in vitro* assays show that the RNAP II CTD and ANP32 jointly enhance cRNA synthesis. The data support a model in which the CTD binds at the interface between FluPol_(E)_ and ANP32, thereby forming a FluPol_(R)_-ANP32A-FluPol_(E)_ complex in which both the FluPol_(R)_ and FluPol_(E)_ are bound to the CTD (**Fig. 5K, e-f**). The CTD is a low complexity disordered region and there is evidence that it drives phase separation and the formation of biomolecular condensates that concentrate RNAP II and transcription regulatory factors (Cramer, 2019). In infected cells, the RNPA II condensates may in addition concentrate the FluPol and its viral and cellular partners, allowing CTD-bound FluPols to switch efficiently between transcription or replication depending on the relative availability of, e.g. 5’ capped RNAs, NP and/or apo FluPol (**Fig. 5K, g**), and perform both activities in a coordinated manner.

Despite many attempts we were unable to provide structural evidence for a ANP32A-CTD-FluPol_(E)_ complex. A possible explanation is that such complexes are stably assembled only when the FluPol_(R)_ is in the context of a vRNP, or in the presence of additional viral (Newcomb et al., 2009) and/or cellular factor(s). Another, non-exclusive explanation, could relate to the fact the CTD-mimicking peptides used for *in vitro* experiments are six repeats in length whereas the full-length CTD consists of 52 repeats in mammals. Moreover, the observed FluPol *de novo* initiation at position 2 of the vRNA (Drncova et al., unpublished) may result in a hook structure at the 5’ end of the nascent cRNA that is less stable and therefore less likely to be bound by a FluPol_(E)_ moiety, which could explain why the asymmetric FluPol_(R)_-FluPol_(E)_ dimer is not observed.

Although a structure for the asymmetric FluPol_(R)_-ANP32-FluPol_(E)_ replicase-encapsidase dimer has so far been obtained only for vRNA-bound FluPol_(R)_, it is generally thought that the same type of asymmetric dimer is formed by a cRNA-bound FluPol_(R)_ (Zhu et al., 2023). Therefore, the model proposed for incoming vRNPs (**Fig. 5K**) can be extended to neo-synthesised, replicating cRNPs and vRNPs, consistent with vRNA synthesis being detected mostly in the chromatin-associated fractions (López-Turiso et al., 1990; Takizawa et al., 2006). In our model, the FluPol_(R)_-associated RNPs could remain anchored to the same CTD repeats and undergo several rounds of cRNA->vRNA or vRNA->cRNA synthesis, while the FluPol_(E)_-associated RNPs could form novel CTD-associated FluPol_(R)_ or FluPol_(T)_ after RNA synthesis is complete. Our model that the RNAP II CTD provides a recruitment platform for the different types of monomeric and multimeric polymerases raises many questions for future studies. What is the sequence of events leading to CTD-anchored replicase complexes? Are the CTD and/or host factors recruited by the CTD controlling the balance between transcription and replication? Are transcribing and replicating FluPol complexes localised to the same RNAP II condensates? Could the viral-induced degradation of RNAP II observed at late time-points during influenza virus infection (Rodriguez et al., 2007; Vreede et al., 2010) favour the release of neo-synthesised vRNPs from the chromatin and their nuclear exportFully addressing these questions will require to overcome technical challenges such as structural analyses on the FluPol in the context of vRNPs and cRNPs, or high-resolution imaging in live-infected cells to visualise the subnuclear localisation of FluPol transcription and replication.

## Material and Methods

### Cells

HEK-293T cells (ATCC CRL-3216) were grown in complete Dulbecco’s modified Eagle’s medium (DMEM, Gibco) supplemented with 10% fetal bovine serum (FBS) and 1% penicillin-streptomycin (Gibco). MDCK cells (provided by the National Influenza Center, Paris, France) were grown in Modified Eagle’s medium (MEM, Gibco) supplemented with 5% FBS and 1% penicillin-streptomycin.

### Plasmids

The A/WSN/33 (WSN) pcDNA3.1-PB2, -PB1, -PA and pCI-NP plasmids were described previously (Krischuns et al., 2022; Lukarska et al., 2017). The A/Anhui/1/2013 (Anhui-H7N9) pCAGGS-PB2, - PB1, -PA and NP plasmids were described previously (Mänz et al., 2016) and were kindly provided by R. Fouchier (Erasmus Medical Center, Netherlands). The B/Memphis/13/2003 pcDNA3.1-PB2, PA plasmids were described previously (Guilligay et al., 2008). The WSN reverse genetics plasmids were kindly provided by G. Brownlee (Sir William Dunn School of Pathology, Oxford, UK) (Fodor et al., 1999). For vRNP reconstitution assays, a pPolI-Firefly plasmid encoding the Firefly luciferase sequence in negative polarity flanked by the 5’ and 3’ non-coding regions of the IAV NS segment was used and the pTK-Renilla plasmid (Promega) was used as an internal control. For RNA quantifications in vRNP reconstitution assays by strand-specific qRT-PCR a pPR7-WSN-NA reverse genetic plasmid was used (Vieira Machado et al., 2006). The WSN and Anhui-H7N9 pCI-PB1-G1 and pCI-PB1-G2 plasmids used for *G. princeps* split-luciferase-based protein-protein complementation assays were constructed as described previously (Cassonnet et al., 2011). The Anhui-H7N9 PA open-reading frame (ORF) was subcloned into the pCI vector and the PA-X ORF was deleted when indicated by introducing silent mutations at the site of ribosomal frameshifting (Jagger et al., 2012). For CTD-binding assays, a C-terminal stretch of RPB1 (108 amino acids) in conjunction with 14 CTD repeats were tagged C-terminally with the *G. princeps* fragments as described previously (Morel et al., 2022). For ANP32A-binding assays, ANP32A was tagged C-terminally with the *G. princeps* fragments as described previously (Long et al., 2019). The pCI-mCherry-CTD-WT and mCherry-CTD-S5A plasmids were generated by replacing the G2 sequence in previously described pCI-G2-CTD constructs (Krischuns et al., 2022). ANP32A encoding the amino acids 1-249 (full-length), 1-199 and 1-149 were subcloned into the pCI vector with a C-terminal V5-tag and SV40 nuclear localisation signals (NLS). pCI-G2-DDX5 and -G2-RED were described previously (Fournier et al., 2014; Munier et al., 2013). All mutations were introduced by an adapted QuikChange site-directed mutagenesis (Agilent Technologies) protocol (Zheng et al., 2004). All primer and plasmid sequences are available upon request.

### Generation of HEK-293T ANP32A and ANP32B knockout cells

CRISPR-Cas9-mediated HEK-293T ANP32A and ANP32B knockout cells (HEK-293T ANP32AB KO) and a control cell line (HEK-293T CTRL) were generated by lentiviral transduction. Guide RNAs (gRNAs) targeting ANP32A (5’-ACCGCTTTGGTAAGTTTGCGATTG-3’ and 5’-AAACCAATCGCAAACTTACCAAAG-3’), ANP32B (5’-ACCGGGAGCTGAGGAACCGGACCC-3’ and 5’-AAACGGGTCCGGTTCCTCAGCTCC-3’) and a non-targeting control (5’-ACCGGTATTACTGATATTGGTGGG-3’ and 5’-AAACCCCACCAATATCAGTAATAC-3’) were annealed and cloned into the BsmBI site of pSicoR-CRISPR-PuroR (a kind gift from Robert Jan Lebbink, (van Diemen et al., 2016)). Replication-incompetent lentiviral particles were generated by transient transfection of HEK-293T cells with the pSicoR-CRISPR-PuroR plasmid harbouring gRNAs targeting either ANP32A, ANP32B or a non-targeting control gRNA as well as with the packaging plasmids pMD2.G and psPAX2 (gifts from Didier Trono, Addgene plasmids # 12259 and # 12260, respectively). Supernatants were harvested 72 hpt, centrifuged to remove cellular debris and passed through 0.45 µm sterile filters. Fresh HEK-293T cells were plated in 6 well plates precoated with poly-D-lysine and were infected with 3 ml of lentiviral supernatant by constant centrifugation at 2.000 rpm for 40 min in the presence of 10 µg/ml Polybrene (Sigma-Aldrich). At 72 hpi, the medium was exchanged to selection medium containing 1 µg/ml puromycin (PAA Laboratories) for one week. Surviving cells were then cloned by limiting dilution in 96-well plates. Individual cell clones were grown and ANP32A and ANP32B protein levels in cellular lysates of individual clones was assessed by immunoblotting. To generate HEK293T cells with a double KO for ANP32A and ANP32B, two rounds of targeting were performed.

### Protein complementation assays

HEK-293T cells were seeded in 96-well white plates (Greiner Bio-One) the day before transfection using polyethyleneimine (PEI-max, #24765-1 Polysciences Inc). Cells were lysed 20-24 hpt in *Renilla* lysis buffer (Promega) for 45 min at room temperature (RT) under steady shaking. *G. princeps* luciferase enzymatic activity due to luciferase reconstitution was measured on a Centro XS LB960 microplate luminometer (Berthold Technologies) using a reading time of 10 s after injection of 50 µl *Renilla* luciferase reagent (Promega). Mean relative light units (RLUs) of technical triplicates of *G. princeps* split-luciferase-based protein-protein complementation assays are represented. In the graphs each dot represents an independently performed biological replicate while at least three experiments were performed in each case. For FluPol-binding assays cells were always co-transfected with plasmids encoding the viral polymerase subunits (PB2, PB1, PA) tagged to one part of the *G. princeps* luciferase and the target protein tagged to the other part. The following combinations of *G. princeps* tags were used while the naming order indicates C- or N-terminal tagging: PB1-G2 and CTD-G1, PB1-G1 and CTD-G2, PB1-G2 and ANP32A-G1, PB1-G1 and ANP32A-G2, PB2-G1 and G2-RED, PB2-G1 and G2-DDX5. For FluPol dimerisation PB2, PA, PB1-G1 and PB1-G2 were co-transfected as described previously (Chen et al., 2019) and when indicated an expression plasmid for mCherry, mCherry-CTD-WT or mCherry-CTD-S5A was added while the total amount of transfected plasmid was adjusted using an empty control plasmid.

### vRNP reconstitution assays

HEK-293T cells were seeded in 96-well white plates (Greiner Bio-One) the day before transfection. Cells were co-transfected with plasmids encoding the vRNP protein components (PB2, PB1, PA, NP), a pPolI-Firefly plasmid encoding a negative-sense viral-like RNA expressing the Firefly luciferase and the pTK-Renilla plasmid (Promega) as an internal control. For FluPol trans-complementation assays, equal amounts of expression plasmids expressing the trans-complementing PA mutants were co-transfected. The fold-change shown for FluPol trans-complementation assays represents the increase above the integrated background of the respective FluPol mutants alone. For FluPol activity rescue experiments in HEK-293T ANP32AB KO cells, expression plasmids encoding FL or truncated versions of ANP32A were co-transfected. Firefly luciferase activity due to viral polymerase activity and Renilla luciferase activity due to RNAP II activity were measured using the Dual-Glo Luciferase Assay system (Promega) according to the manufacturer’s instructions. The graphs represent viral polymerase activity normalised to RNAP II activity. Luciferase activities were measured in technical duplicates at 20-24 hpt or at 48 hpt when indicated. In the graphs each dot represents an independently performed biological replicate while at least three experiments were performed in each case. For the quantification of mRNA, cRNA and vRNA levels, HEK-293T cells were seeded in 12-well plates and transfected with plasmids encoding the vRNP protein components (PB2, PB1, PA, NP) and a low amount of WSN-NA RNA expressing plasmid (5 ng/well). Total RNA was isolated at 24 hpt with RNeasy Mini columns according to the manufacturer’s instructions (RNeasy Kits, Qiagen) and strand-specific RT-qPCRs were performed as described previously (Kawakami et al., 2011). Briefly, total RNA was reverse transcribed using primers specific for NA mRNA, cRNA, vRNA and, when indicated, the cellular glyceraldehyde 3-phosphate dehydrogenase (GADPH) with SuperScript^TM^ III Reverse Transcriptase (Invitrogen) and quantified using SYBR-Green (Roche) with the LightCycler 480 system® (Roche). RNA levels were normalised to GAPDH when indicated and analysed using the 2^−ΔΔCT^ as described before (Livak and Schmittgen, 2001).

### Antibodies and immunoblots

Total cell lysates were prepared in RIPA cell lysis buffer as described before (Krischuns et al., 2018). Proteins were separated by SDS-PAGE using NuPAGE™ 4-12% Bis-Tris gels (Invitrogen) and transferred to nitrocellulose membranes which were incubated with primary antibodies directed against pS5 CTD (clone 3E8), mCherry (26765-1-AP), V5 (SV5-Pk1), Tubulin (B-5-1-2), PA (Da Costa et al., 2015), PB2 (GTX125925, GeneTex), ANP32A (AV40203, Sigma), ANP32B (EPR14588) and subsequently with HRP-tagged secondary antibodies (Jackson Immunoresearch). Membranes were revealed with the ECL2 substrate according to the manufacturer’s instructions (Pierce) and chemiluminescence signals were acquired using the ChemiDoc imaging system (Bio-Rad) and analysed with ImageLab (Bio-Rad).

### Production and characterisation of recombinant viruses

The recombinant PA mutant influenza viruses (PA K289A, R454A, K635A, R638A) used for serial cell culture passaging were generated by reverse genetics followed by plaque purification under agarose overlay and one round of viral amplification (P1) as described previously (Lukarska et al., 2017). For the serial passaging MDCK cell were infected at a multiplicity of infection (MOI) of 0.0001 and incubated for 3 days at 37°C in DMEM containing TPCK-Trypsin (Sigma) at a final concentration of 1 µg/mL. Between each passage, viral supernatants were titrated on MDCK cells in a plaque assay as described before (Matrosovich et al., 2006). Viral RNA was extracted from 140 µL of viral stocks using the QIAamp Viral RNA Mini kit (Qiagen) according to the manufacturer’s instructions. For next-generation sequencing of the full viral genome reverse transcription and amplification of the eight genomic segments were performed using the RT-qPCR protocol adapted by the National Influenza Center (Institut Pasteur) (Watson et al., 2013) as described previously (Chen et al., 2019). Next-generation sequencing was performed by the P2M facility at Institut Pasteur using the Nextera XT DNA Library Preparation kit (Illumina), the NextSeq 500 sequencing systems (Illumina) and the CLC Genomics Workbench 9 software (Qiagen) for analysis. Recombinant viruses with a selected subset of observed second-site mutations were generated by reverse genetics as described previously sja(Sjaastad et al., 2018). In brief, HEK-293T cells were seeded in 6-well plates the day before transfection with the WSN reverse genetics plasmids in Opti-MEM (Gibco) using FuGene®6 (Promega) according to the manufacturer’s instructions (Fodor et al., 1999). The next day, MDCK were added to the wells in 500 µL Opti-MEM containing 0.5 µg/mL TPCK trypsin (Sigma). The following days, 500 µL Opti-MEM containing 1 and 2 µg/mL TPCK trypsin, respectively, were added. One day later, the supernatants were harvested, centrifuged and stored at −80°C. The reverse genetics supernatants were titrated on MDCK cells and plaque diameters were measured upon staining with crystal violet using Fiji (Schindelin et al., 2012).

### Influenza virus polymerase Zhejiang-H7N9 (WT, 4M, PA K289A+C489R)

The previously described and cloned pFastBac Dual vector encoding for the A/Zhejiang/DTID-ZJU01/2013 (H7N9) polymerase heterotrimer subunits, PA (Uniprot: M9TI86), PB1 (Uniprot: M9TLW3), and PB2 (Uniprot: X5F427) was used as a starting point (Zhejiang-H7N9 WT) (Kouba et al., 2023). The mutations PA E349K, PA R490I, PB1 K577G and PB2 G74R were introduced by a combination of PCRs and Gibson assembly (Zhejiang-H7N9 4M). A similar approach has been applied to clone the FluPol Zhejiang-H7N9 PA K289A+C489R mutant. Plasmid sequences were confirmed by sanger sequencing for each polymerase subunit. All primer and plasmid sequences are available upon request.

FluPols Zhejiang-H7N9 (WT / 4M / PA K289A+C489R) were produced using the baculovirus expression system in *Trichoplusia ni* High 5 cells. For large-scale expression, cells at 0.8-1E06 cells/mL concentration were infected by adding 1% of virus. Expression was stopped 72 to 96 h after the day of proliferation arrest and cells were harvested by centrifugation (1.000g, 20 min at 4°C).

Cells were disrupted by sonication for 4 min (5 s ON, 20 s OFF, 40% amplitude) on ice in lysis buffer (50 mM HEPES pH 8, 500 mM NaCl, 0.5 mM TCEP, 10% glycerol) with cOmplete EDTA-free Protease Inhibitor Cocktail (Roche). After lysate centrifugation at 48.384g during 45 min at 4°C, ammonium sulfate was added to the supernatant at 0.5 g/mL final concentration. The recombinant protein was then collected by centrifugation (45 min, 4°C at 70.000g), re-suspended in the lysis buffer, and the procedure was repeated another time. FluPol Zhejiang-H7N9 was then purified using His60 NiNTA Superflow resin (Takara Bio) from the soluble fraction. Bound proteins were subjected to two sequential washes steps using (i) the lysis buffer supplemented by 1 M NaCl and (ii) the lysis buffer supplemented by 50 mM imidazole. Remaining bound proteins were eluted using the lysis buffer supplemented by 500 mM imidazole. Fractions with FluPol Zhejiang-H7N9 were pooled and directly subjected to a strep-tactin affinity purification (IBA, Superflow). Bound proteins were eluted using the lysis buffer supplemented by 2.5 mM d-desthiobiotin and protein-containing fractions were pooled and diluted with an equal volume of buffer (50 mM HEPES pH 8, 0.5 mM TCEP, 10% glycerol) before loading on a third affinity column HiTrap Heparin HP 5 mL (Cytiva). A continuous gradient of lysis buffer supplemented with 1 M NaCl was applied over 15 CV, and FluPol Zhejiang-H7N9 was eluted as single species at ∼800 mM NaCl. Pure and acid nucleic free FluPol Zhejiang-H7N9 was dialysed overnight in a final buffer (50 mM HEPES pH 8, 500 mM NaCl, 2 mM TCEP, 5% glycerol), concentrated with Amicon Ultra-15 (50 kDa cutoff), flash-frozen and stored at −80°C for further use.

### Influenza virus nucleoprotein Anhui-H7N9 monomeric mutant (R416A)

The influenza virus nucleoprotein (NP) Anhui-H7N9 gene was amplified from a pCAGGS plasmid (gift from Ron Fouchier) (Mänz et al., 2016) and introduced by Gibson assembly in a pLIB vector with an N-terminal double strep-tag followed by a human rhinovirus (HRV) 3C protease cleavage site. The NP R416A mutation was introduced by a combination of PCRs and Gibson assembly. Introduction of the desired mutation was confirmed by sanger sequencing. Monomeric influenza virus Anhui-H7N9 NP R416A was produced using the baculovirus expression system in *Trichoplusia ni* High 5 cells. For large-scale expression, cells at 0.8-1E06 cells/mL concentration were infected by adding 1% of virus. Expression was stopped 72 to 96 h after the day of proliferation arrest and cells were harvested by centrifugation (1.000g, 20 min at 4°C).

Cells were disrupted by sonication for 4 min (5 s ON, 15 s OFF, 40% amplitude) on ice in lysis buffer (50 mM HEPES pH 8, 500 mM NaCl, 2 mM TCEP, 5% glycerol) with cOmplete EDTA-free Protease Inhibitor Cocktail (Roche). After lysate centrifugation at 48.384g for 45 min at 4°C, the soluble fraction was loaded on a StrepTrap HP 5 mL (Cytiva). Bound proteins were eluted using the lysis buffer supplemented by 2.5 mM d-desthiobiotin. Protein-containing fractions were pooled and diluted with an equal volume of buffer (50 mM HEPES pH 8, 2 mM TCEP, 5% glycerol) before loading on an affinity column HiTrap Heparin HP 5 mL (Cytiva). A continuous gradient of lysis buffer supplemented with 1 M NaCl was applied over 15 CV, and Anhui-H7N9 NP R416A was eluted as single species at ∼500 mM NaCl without nucleic acids (A_260/280_: 0.6). Pure and acid nucleic free Anhui-H7N9 NP R416A was dialysed overnight in a final buffer (50 mM HEPES pH 8, 300 mM NaCl, 1 mM TCEP, 5% glycerol) together with N-terminal his-tagged HRV 3C protease (ratio 1:5 w/w). Tag-cleaved Anhui-H7N9 NP R416A was subjected to a last Ni-sepharose affinity to remove the HRV 3C protease, further concentrated with Amicon Ultra-15 (10 kDa cutoff), flash-frozen and stored at −80°C for later use.

### Acidic Nuclear Phosphoprotein 32A

Human and chicken ANP32A genes (GeneScript) were introduced in a pETM11 vector with an N-terminal 6xHis-tag followed by a Tobacco Etch Virus (TEV) protease cleavage site. ANP32A constructs were expressed in BL21(DE3) *E.coli* cells. Expression was induced when absorbance reached 0.6, with 1 mM IPTG, incubated for 4 h at 37°C. Cells were harvested by centrifugation (1.000g, 20 min at 4°C).

Cells were disrupted by sonication for 5 min (5 s ON, 15 s OFF, 50% amplitude) on ice in lysis buffer (50 mM HEPES pH 8, 150 mM NaCl, 5 mM beta-mercaptoethanol (BME)) with cOmplete EDTA-free Protease Inhibitor Cocktail (Roche). After lysate centrifugation at 48.384g for 45 min at 4°C, the soluble fraction was loaded on a HisTrap HP 5 mL column (Cytiva). Bound proteins were subjected to a wash step using the lysis buffer supplemented by 50 mM imidazole. Remaining bound proteins were eluted using the lysis buffer supplemented by 500 mM imidazole. Fractions containing ANP32A were dialysed overnight in the lysis buffer (50 mM HEPES pH 8, 150 mM NaCl, 5 mM BME) together with N-terminal his-tagged TEV protease (ratio 1:5 w/w). Tag-cleaved ANP32A protein was subjected to a Ni-sepharose affinity column to remove the TEV protease, further concentrated with Amicon Ultra-15 (3 kDa cutoff) and subjected to a Size-Exclusion Chromatography using a Superdex 200 Increase 10/300 GL column (Cytiva) in a final buffer containing 50 mM HEPES pH 8, 150 mM NaCl, 2 mM TCEP. Fractions containing exclusively ANP32A were concentrated with Amicon Ultra-15 (3 kDa cutoff), flash-frozen and stored at −80°C for later use.

Human ANP32A truncation constructs (1-199 and 144-249) were generated, expressed and protein purified as previously described (Camacho-Zarco et al., 2020). The human ANP32A 1-149 construct was a gift from Cynthia Wolberger (Addgene plasmid # 67241, (Huyton and Wolberger, 2007)) and was expressed and protein purified as previously described (Camacho-Zarco et al., 2020).

### *De novo* FluPol replication activity

Synthetic vRNA loop (“v51_mut_S”) (5’-pAGU AGA AAC AAG GGU GUA UUU UCC CCU CUU UUU GUU UCC CCU GCU UUU GCU −3’) (IDT) was used for all *in vitro* replication activity assays. For all *de novo* replication activity assays, 2.4 µM FluPol Zhejiang-H7N9 (WT or 4M) were mixed with (i) 0.8 µM v51_mut_S and/or (ii) 8 µM ANP32A and/or (iii) 16 µM pS5 CTD(6mer) (respective molar ratio: 3 FluPols: 1 template: 10 ANP32: 20 pS5 CTD). Reactions were launched at 30°C for 4 h by adding ATP/GTP/CTP/UTP (AGCU) or only ATP/GTP/CTP (AGC) at 100 µM/NTP, 0.75 µCi/ml α-32P ATP and MgCl_2_ in a final assay buffer containing 50 mM HEPES pH 8, 150 mM NaCl, 2 mM TCEP, 100 µM/NTP, 1 mM MgCl_2_. Reactions were stopped by adding 2X RNA loading dye, heating 5 min at 95°C and immediately loaded on a 20% TBE-7M urea-polyacrylamide gel. Each gel was exposed on a storage phosphor screen and read with an Amersham Typhoon scanner (Cytiva). For each gel the decade markers system (Ambion) was used.

### Next Generation Sequencing of *in vitro* FluPol replication products

To confirm the identity of the replication products (full-length cRNA product and stalled cRNA product) and address the discrepancy between expected size and urea-PAGE migration, sequencing of the total reaction product was performed from a *de novo* FluPol replication assay. 2.4 µM FluPol Zhejiang-H7N9 4M were mixed with (i) 0.8 µM v51_mut_S, (ii) 8 µM ANP32A and (iii) 16 µM pS5 CTD(6mer) (respective molar ratio: 3 FluPols: 1 template: 10 ANP32A: 20 pS5 CTD). Reaction was launched at 30°C for 4 h by adding ATP/GTP/CTP/UTP at 100 µM/NTP and MgCl_2_ in a final assay buffer containing 50 mM HEPES pH 8, 150 mM NaCl, 2 mM TCEP, 100 µM/NTP, 1 mM MgCl_2_. After completion of the reaction, proteins were removed using the Monarch RNA Cleanup Kit (NEB) and 5’-mono-phosphorylated v51_mut_S templates were specifically digested with a terminator 5’-phosphate-dependent exonuclease (Biosearch technologies) and subjected again to the Monarch RNA Cleanup Kit (NEB). Total remaining RNAs were used for next-generation sequencing.

Sample concentration and fragment size distribution were checked with the RNA Pico assay (Bioanalyzer, Agilent). 6 ng of sample was treated with 2 units T4 Polynucleotide Kinase (NEB) and incubated at 37°C for 30 min, followed by heat inactivation at 65°C for 20 min. The treated samples were then taken into library preparation following the Takara SMARTer small RNA sequencing kit according to the manufacturer’s instructions with 13 cycles of PCR. The final library fragment size distribution was checked with the High Sensitivity Bioanalyzer assay. 8 pM of library was sequenced on a MiSeq to generate 76 bp single-end reads.

Resulting total reads (13.073.371) were trimmed using Cutadapt v2.3 (Kechin et al., 2017) with settings recommended by the library preparation kit (-u 3 -a ‘AAAAAAAAAA’) omitting the filter for only reads longer than 20 bases (10.991.864). Trimmed reads were subsequently used as an input for pattern lookup performed using Python scripts (**Source data file 1**). All reads exactly starting by these motifs have been kept for further analysis: “AGCAAAAGCA”/“GCAAAAGCA”/“CAAAAGCA”/“AAAAGCA” (7.776.959 reads). Each read starting exactly by these motifs “5’-AGCAAAAGCAGGGG” / “5’ -GCAAAAGCAGGGG” / “5’-CAAAAGCAGGGG” / “5’ -AAAAGCAGGGG” were counted and plotted (**Fig. 3.H**). Each read matching the exact sequence of a full-length replication product starting by “5’-AGC AAA…ACU-3’” (51-mer), “5’-_GC AAA… ACU-3’” (50-mer), “5’-C AAA… ACU-3’” (49-mer) or “5’-AAA… ACU-3’” (48-mer) were counted and plotted as percentage (**Figure 3.I**).

### Electron microscopy

D*e novo* replication states were trapped by mixing 2.4 µM FluPol Zhejiang-H7N9 4M with (i) 0.8 µM v51_mut_S, (ii) 8 µM chANP32A, (iii) 16 µM pS5 CTD(6mer) and (iv) 4 µM Anhui-H7N9 NP R416A in a final buffer containing 50 mM HEPES pH 8, 150 mM NaCl, 2 mM TCEP, 1 mM MgCl_2_, ATP/GTP/CTP at 100 µM/NTP, and 100 µM of non-hydrolysable UpNHpp (Jena Bioscience). The *de novo* reaction mix was incubated for 4 h at 30°C. Before proceeding to grids freezing, the sample was centrifuged for 5 min at 11.000g and stored at 4°C before proceeding to grids freezing. For grids preparation, 1.5 µl of sample was applied on each sides of plasma cleaned (Fischione 1070 Plasma Cleaner: 1 min 10 s, 90% oxygen, 10% argon) grids (UltrAufoil 1.2/1.3, Au 300). Excess solution was blotted for 3 to 5 s, blot force 0, 100% humidity, at 4°C, with a Vitrobot Mark IV (ThermoFisher) before plunge-freezing in liquid ethane. Automated data collection was performed on a TEM Titan Krios G3 (Thermo Fisher) operated at 300 kV equipped with a K3 (Gatan) direct electron detector camera and a BioQuantum energy filter, using EPU. Coma and astigmatism correction were performed on a carbon grid. Micrographs were recorded in counting mode at a ×105,000 magnification giving a pixel size of 0.84 Å with defocus ranging from −0.8 to −2.0 µm. Gain-normalised movies of 40 frames were collected with a total exposure of ∼40 e^−^/Å^2^.

For Zhejiang-H7N9 PA K289A+C489R structures, 0.8 µM FluPol Zhejiang-H7N9 PA K289A+C489R was mixed with (i) 0.8 µM v51_mut_S and 16 µM pS5 CTD(6mer) in a final buffer containing 50 mM HEPES pH 8, 150 mM NaCl, 2 mM TCEP. After 1 h incubation on ice, the sample was centrifuged for 5 min, 11.000g and stored at 4°C before proceeding to grids freezing. For grid preparation, 1.5 µl of sample was applied on each sides of plasma cleaned (Fischione 1070 Plasma Cleaner: 1 min 10 s, 90% oxygen, 10% argon) grids (UltrAufoil 1.2/1.3, Au 300). Excess solution was blotted for 3 to 5 s, blot force 0, 100% humidity, at 4°C, with a Vitrobot Mark IV (ThermoFisher) before plunge-freezing in liquid ethane. Automated data collection was performed on a TEM Glacios (ThermoFisher) operated at 200 kV equipped with a F4i (ThermoFisher) direct electron detector camera and a SelectrisX energy filter, using EPU. Coma and astigmatism correction were performed on a carbon grid. Micrographs were recorded in counting mode at a ×130,000 magnification giving a pixel size of 0.907 Å with defocus ranging from −0.8 to −2.0 µm. EER movies were collected with a total exposure of ∼40 e^−^/Å^2^.

### Image processing

For the TEM Titan Krios dataset, movie drift correction was performed using Relion’s Motioncor implementation, with 7×5 patch, using all movie frames (Zheng et al., 2017). All additional initial image processing steps were performed in cryoSPARC v3.3 (Punjani et al., 2017). CTF parameters were determined using “Patch CTF estimation”, realigned micrographs were then manually inspected and low-quality images were manually discarded. To obtain an initial 3D reconstruction of FluPol Zhejiang-H7N9 4M, particles were automatically picked on few hundreds micrographs using a circular blob with a diameter ranging from 100 to 140 Å. Particles were extracted using a box size of 360 x 360 pixels^2^, 2D classified and subjected to an “ab-initio reconstruction” job. The best initial model was further used to prepare 2D templates. Template picking was then performed using a particle diameter of 120 Å and particles extracted from dose-weighted micrographs. Successive 2D classifications were used to eliminate particles displaying poor structural features. All remaining particles were then transferred to Relion 4.0. Particles were divided in subset of 300k to 500k particles and subjected to multiple 3D classification with coarse image-alignment sampling using a circular mask of 180 Å. For each similar FluPol conformation, particles were grouped and subjected to 3D masked refinement followed by multiple 3D classification without alignment or using local angular searches. Once particles properly classified, bayesian polishing was performed and re-extracted “shiny” particles were subjected to a last 3D masked refinement. Post-processing was performed in Relion using an automatically or manually determined B-factor. For each final map, reported global resolution is based on the FSC 0.143 cut-off criteria. Local resolution variations were estimated in Relion. Detailed image processing information is shown in Fig. S7.

For the TEM Glacios dataset, EER fractionation was set to 24, giving ∼1e^−^/Å^2^ per resulting fraction. Movie drift correction was performed using Relion’s Motioncor implementation, with 5×5 patch (Zheng et al., 2017). All additional initial image processing steps were performed in cryoSPARC v4.1 (Punjani et al., 2017). CTF parameters were determined using “Patch CTF estimation”, realigned micrographs were then manually inspected and low-quality images were manually discarded. To obtain an initial 3D reconstruction of FluPol Zhejiang-H7N9 PA K289A+C489R, particles were automatically picked on few hundreds micrographs using a circular blob with a diameter ranging from 100 to 140 Å. Particles were extracted using a box size of 340 x 340 pixels^2^, 2D classified and subjected to an “ab-initio reconstruction” job. The best initial model was further used to prepare 2D templates. Template picking was then performed using a particle diameter of 120 Å and particles extracted from dose-weighted micrographs. Successive 2D classifications were used to eliminate particles displaying poor structural features. All remaining particles were subjected to an “Heterogenenous refinement” job, with 3 symmetrical dimers and 3 monomers as initial 3D models. Particles assigned to the monomeric 3D classes were subjected to a final non-uniform refinement. Post-processing was performed in Relion using an automatically determined B-factor. Reported global resolution is based on the FSC 0.143 cut-off criteria. Local resolution variations were estimated in Relion. Detailed image processing information is shown in Fig. S9.

### Model building and refinement

Using the FluPol Zhejiang-H7N9 elongation complex as starting point (PDB: 7QTL) (Kouba et al., 2023), atomic models were constructed by iterative rounds of manual model building with COOT and real-space refinement using Phenix (Afonine et al., 2018). Validation was performed using Phenix. Model resolution according to the cryo-EM map was estimated at the 0.5 FSC cutoff. Figures were generated using ChimeraX (Goddard et al., 2018).

## Acknowledgments

We thank Ron Fouchier (Erasmus MC Department of Viroscience, Netherlands), George Brownlee (Oxford University), Bernard Delmas (INRAE, France), Robert-Jan Lebbink (Utrecht University, Netherlands) and Sandie Munier (Institut Pasteur, France) for sharing plasmids and antibodies. We thank Martin Pelosse for support in using the Eukaryotic Expression Facility at EMBL Grenoble; Aldo R. Camacho-Zarco and Martin Blackledge for providing huANP32A truncated constructs proteins; Daphne Walter and Vladimir Benes from the EMBL GeneCore facility for the next generation sequencing data; Wojtek Galej, Sarah Schneider and Romain Linares for access to the Glacios at EMBL Grenoble; Guy Schoehn and Eleftherios Zarkadas for access to the Glacios at IBS Grenoble; Aymeric Peuch for support with using the joint EMBL-IBS computer clusters. We acknowledge the European Synchrotron Radiation Facility and PSB for access to the Titan Krios CM01, and Michael Hons for assistance with data collection. We thank the Pasteur International Bioresources Network (PIBNet) for genome sequencing of recombinant viruses.

## SUPPLEMENTARY FIGURES

**Figure S1:**
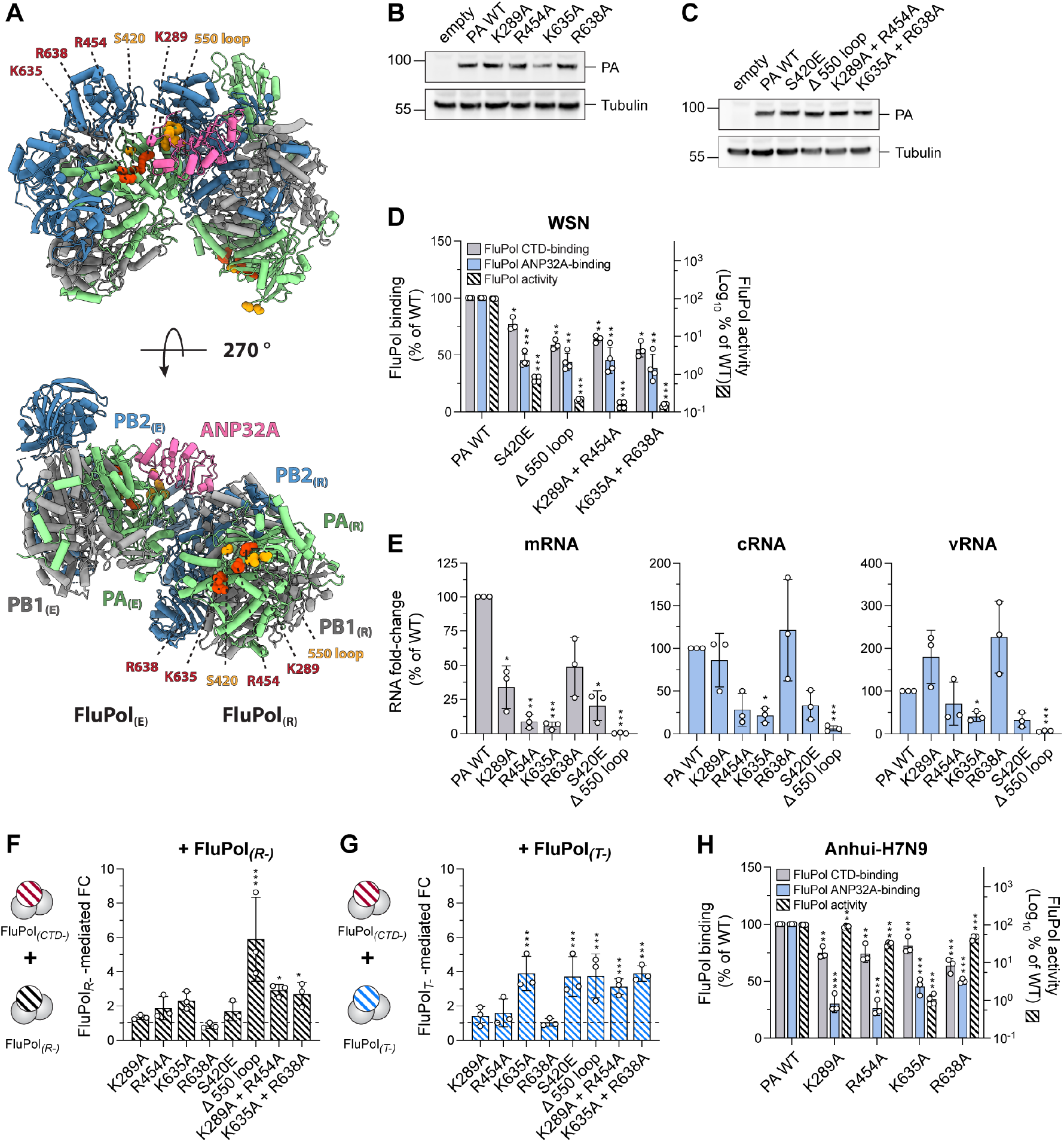
FluPol CTD-binding interface is essential for replication of the viral genome. **(A)** Model of the Flu_A_Pol replication complex (FluPol_(R)_, FluPol_(E)_ and ANP32A) based on a Flu_C_Pol Cryo-EM structure (C/Johannesburg/1/1966 structure, PDB: 6XZR, (Carrique et al., 2020) Ribbon diagram representation with PA in green, PB1 in grey, PB2 in blue and ANP32A in pink. Key FluPol CTD-binding residues PA K289, R454, K635 and R638 are highlighted in red and PA S420 and the PA 550 loop in orange in either the FluPol_(E)_ (top) or FluPol_(R)_ (bottom) conformation. **(B-C)** HEK-293T cells were co-transfected with expression plasmids for WSN PB1, PB2 and PA with the indicated mutation. Cell lysates were analysed by western blot using antibodies specific for PA and tubulin. Uncropped gels are provided as a source data file. **(D)** Cell-based WSN FluPol binding and activity assays. Left Y-axis (linear scale): FluPol binding to the CTD (grey bars) and ANP32A (blue bars) was assessed using split-luciferase-based complementation assays as in Fig. 1C. Luminescence signals due to luciferase reconstitution are represented as a percentage of PA WT. Right Y-axis (logarithmic scale): FluPol PA CTD-binding mutant activity (hatched bars) was measured by vRNP reconstitution in HEK-293T cells as in Fig. 1C. Luminescence signals are represented as a percentage of PA WT. *p < 0.033, **p < 0.002, ***p < 0.001 (one-way ANOVA; Dunnett’s multiple comparisons test). **(E)** WSN vRNPs with the indicated PA mutations were reconstituted in HEK-293T cells by transient transfection using a plasmid encoding the NA vRNA segment. Steady-state levels of NA mRNA, cRNA and vRNA were quantified by strand-specific RT-qPCR (Kawakami et al., 2011) and are represented as a percentage of PA WT. *p < 0.033, **p < 0.002, ***p < 0.001 (one-way repeated measure ANOVA; Dunnett’s multiple comparisons test). **(F-G):** FluPol trans-complementation assay. WSN vRNPs with the indicated PA CTD-binding mutations were reconstituted in HEK-293T cells using a model vRNA encoding the Firefly luciferase, and were co-expressed with (F) a replication-defective (FluPol_(R-)_, PA K664M) or (G) a transcription-defective (FluPol_(T-)_, PA D108A, (Dias et al., 2009)) FluPol mutant. Luminescence was measured at 48 hpt and signals were normalised to a transfection control, and are represented as the fold-change (FC) relative to the background. *p < 0.033, **p < 0.002, ***p < 0.001 (two-way ANOVA; Sidak’s multiple comparisons test). **(H)** Cell-based Anhui-H7N9 FluPol binding and activity assays using a plasmid encoding PA in which the PA-X (PA-ΔX) ORF frame was deleted as described previously (Jagger et al., 2012). Left Y-axis (linear scale): FluPol binding to the CTD (grey bars) and ANP32A (blue bars) was assessed using split-luciferase-based complementation assays as in Fig. 1C. Luminescence signals due to luciferase reconstitution are represented as a percentage of PA WT. Right Y-axis (logarithmic scale): FluPol PA CTD-binding mutant activity (hatched bars) was measured by vRNP reconstitution in HEK-293T cells as in Fig. 1C. Luminescence signals are represented as a percentage of PA WT. **p < 0.002, ***p < 0.001 (one-way ANOVA; Dunnett’s multiple comparisons test).

**Figure S2:**
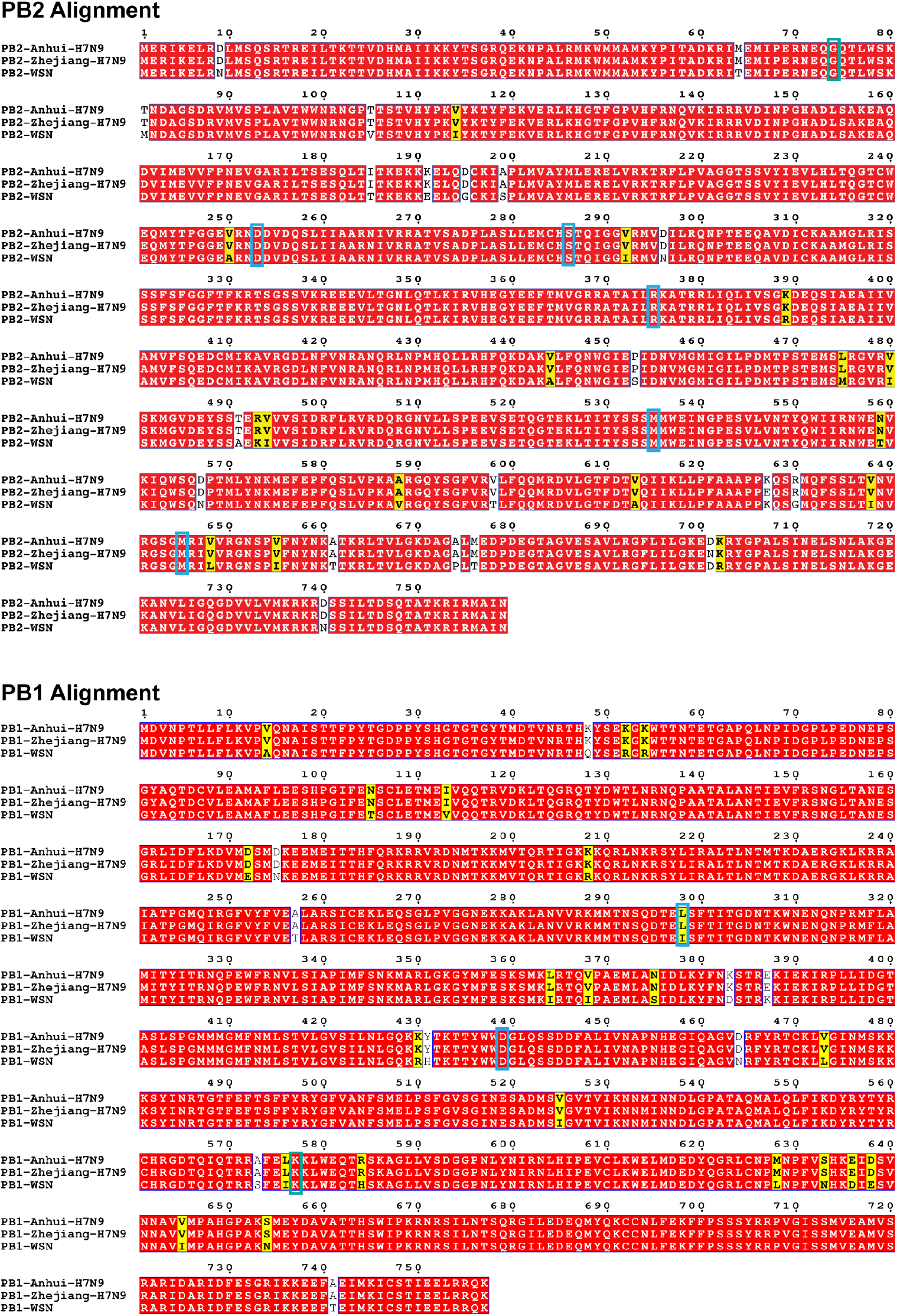
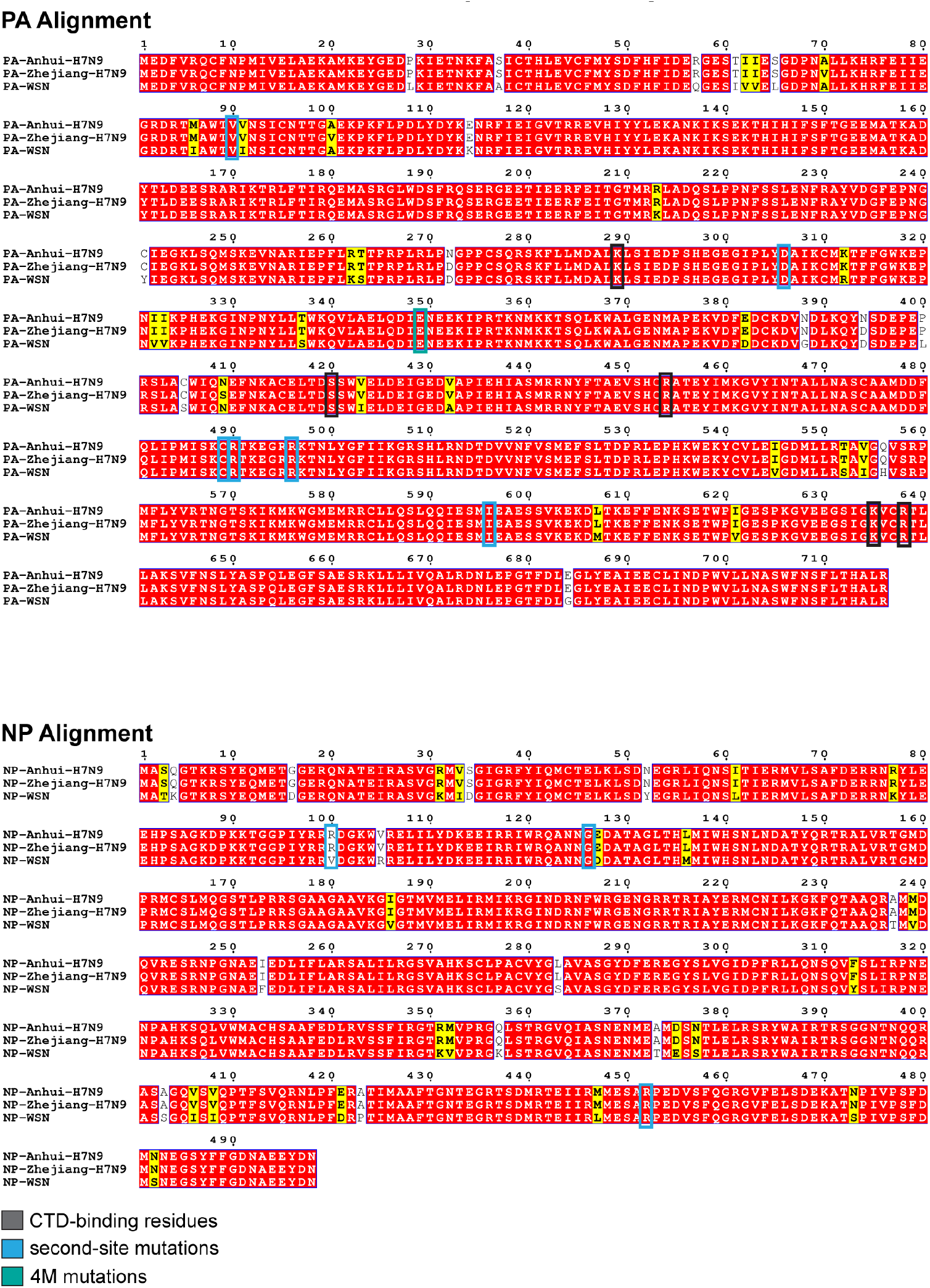
PB2, PB1, PA and NP protein sequence alignments of the influenza A virus strains used in this study. Protein sequences from A/WSN1933(H1N1), A/Anhui/01/2013 (H7N9) and A/Zhejiang/DTID-ZJU01/2013(H7N9) were obtained from UniProt (https://www.uniprot.org/), aligned with CLC Main Workbench 21 and visualised by Espript 3.0 [55]. Identical and similar residues are indicated in red or yellow, respectively. FluPol CTD binding subjected to mutagenesis are indicated by grey boxes, FluPol second-site mutations which appeared upon passaging of FluPol CTD-binding mutant viruses by blue boxes and dimer interrupting FluPol 4M residues (PB2 G74, PA E349, PB1 K577) by green boxes.

**Figure S3:**
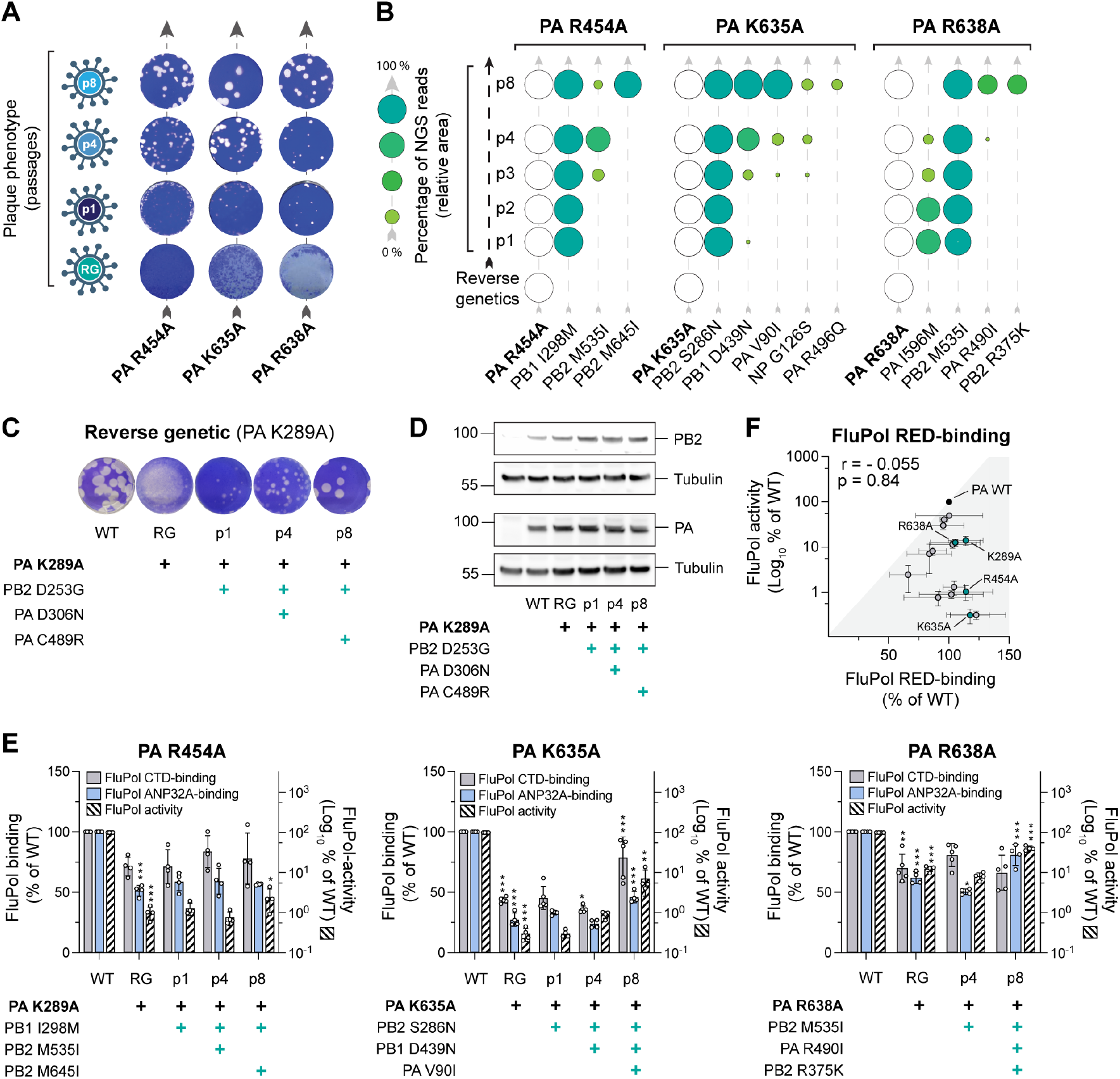
Serial passaging of mutant viruses with mutations at the FluPol-CTD interface selects for adaptive mutations which restore FluPol binding to the CTD and ANP32A. **(A)** Plaque phenotypes of passaged WSN PA R454A, K635A, R638A mutant viruses. Representative images of crystal violet stained plaque assays of the supernatants collected after reverse genetics (RG), p1, p4 and p8 are shown. **(B)** The viral genomes of passaged WSN PA R454A, K635A, R638A mutant viruses were sequenced by short-read NGS after p1 - p4 and p8 (from bottom to top). Initial FluPol CTD-binding mutation (white circles) and second site mutations on PB1, PB2, PA and NP (green circles) are indicated. Only second-site mutations found in ≥ 10% of reads in at least one passage are shown. The fraction of reads showing a given mutation is indicated by the area of the circle, and by different shades of green that represent <25, 25-50, 50-75 and >75 % of reads (schematic on the left). **(C)** Characterisation of recombinant WSN mutant viruses. Recombinant viruses with the primary FluPol mutation PA K289A and the indicated combinations of FluPol second-site mutations (PB2 D253G, PA D306N, PA C489R) were generated by reverse genetics and subjected to plaque assay. Representative images of plaque assays stained by crystal violet are shown. **(D)** HEK-293T cells were co-transfected with expression plasmids for WSN PB1, PB2 and PA with the indicated mutation. Cells lysates were analysed by western blot using antibodies specific for PB2, PA and tubulin. Uncropped gels are provided as a source data file. **(E)** The indicated combinations of WSN FluPol second-site mutations which occurred during passaging of PA R454A, K635A and R638A mutant viruses were characterised in cell-based FluPol binding and activity assays. Left Y-axis (linear scale): FluPol binding to the CTD (grey bars) and ANP32A (blue bars) was assessed using split-luciferase-based complementation assays as in Fig. 1C. Luminescence signals due to luciferase reconstitution are represented as a percentage of FluPol WT. Right Y-axis (logarithmic scale): FluPol activity (hatched bars) was measured by vRNP reconstitution in HEK-293T cells as in Fig. 1C. Luminescence signals are represented as a percentage of FluPol WT. Stars indicate statistical significance compared to the previous passage. *p < 0.033, **p < 0.002, ***p < 0.001 (one-way ANOVA; Tukey’s multiple comparisons test). **(F)** For each initial FluPol mutant (PA K289A, R454A, K635A, R638A, green dots) and FluPol genotype observed during serial passaging (grey dots), FluPol binding to RED (x-axis) was plotted against the FluPol activity as measured in a vRNP reconstitution assay (y-axis). (Pearson Correlation Coefficient (r))

**Figure S4:**
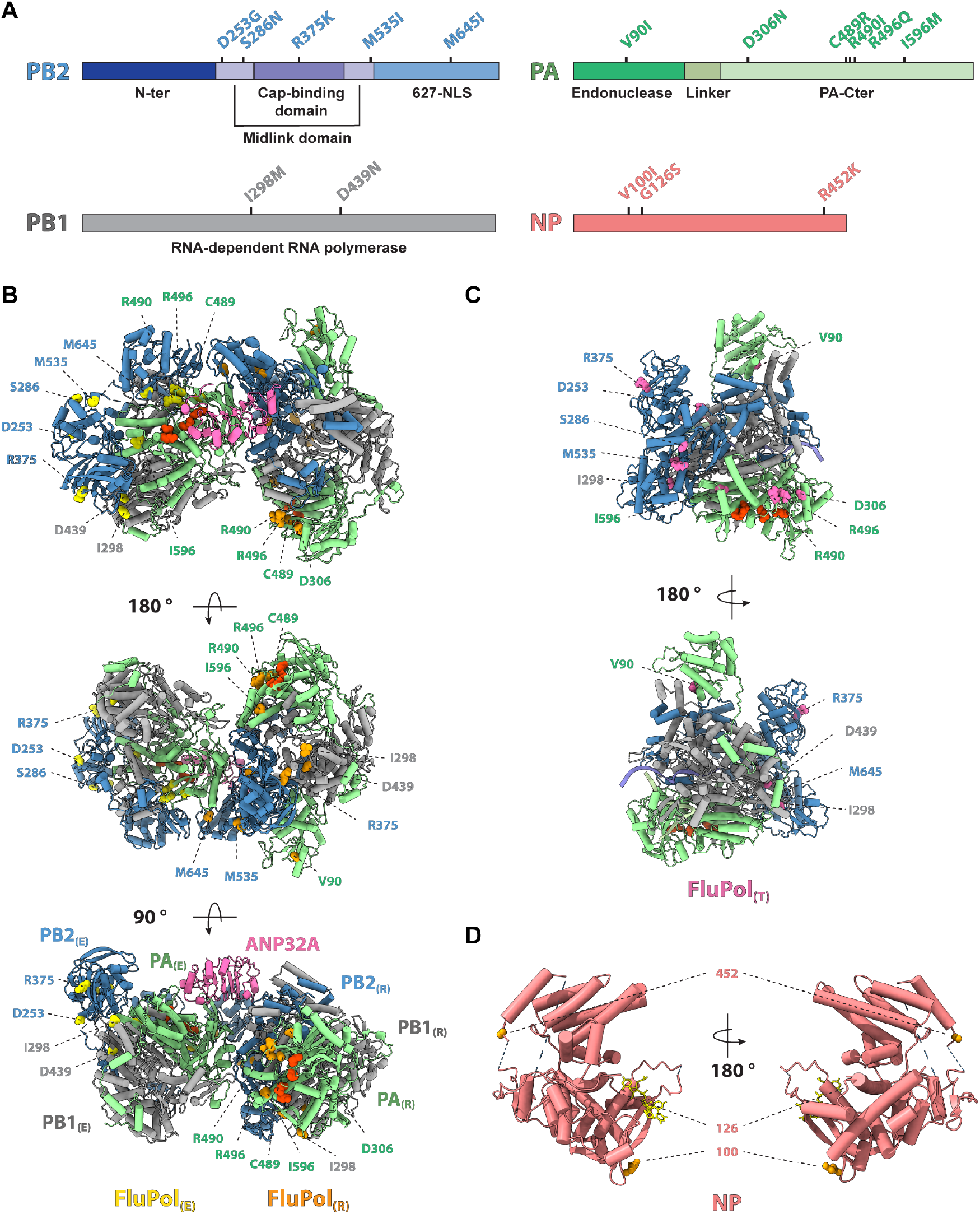
FluPol and NP second-site mutations which were selected during serial cell culture passaging of PA mutant viruses. **(A)** Second-site mutations which were selected on FluPol and NP during serial passaging of recombinant plaque-purified PA mutant (K289A, R454A, K635A, R638A) viruses were mapped on a linear domain representation of PB2, PB1, PA and NP. Only second-site mutations found in ≥ 10% of reads in at least one passage are shown. **(B)** Model of the Flu_A_Pol replication complex (FluPol_(R)_, FluPol_(E)_ and ANP32A) based on a Flu_C_Pol Cryo-EM structure (C/Johannesburg/1/1966 structure, PDB: 6XZR, (Carrique et al., 2020)). Ribbon diagram representation with PA in green, PB1 in grey, PB2 in blue and ANP32A in pink. Initially mutated FluPol PA residues (PA K289A, R454A, K635A and R638A) are highlighted in red and residues corresponding to FluPol second-site mutations were mapped on the FluPol_(R)_ in orange and FluPol_(E)_ in yellow, respectively. **(C)** Representation of the Cryo-EM structure of the FluPol heterotrimer in the FluPol_(T)_ conformation, bound to a 3’5’ vRNA promoter and a short capped RNA primer (A/NT/60/1968, PDB: 6RR7, (Fan et al., 2019)). Ribbon diagram representation with PA in green, PB1 in grey, and PB2 in blue. Initially mutated FluPol PA residues (PA K289A, R454A, K635A and R638A) are highlighted in red and residues corresponding to FluPol second-site mutations in pink. **(D)** Residues corresponding to NP second-site mutations were mapped on the RNA-bound crystal structure of a monomeric NP mutant (PDB: 7DXP, (Tang et al., 2021)). Ribbon diagram representation with NP in salmon, RNA in yellow and residues corresponding to NP second-site mutations in orange.

**Figure S5:**
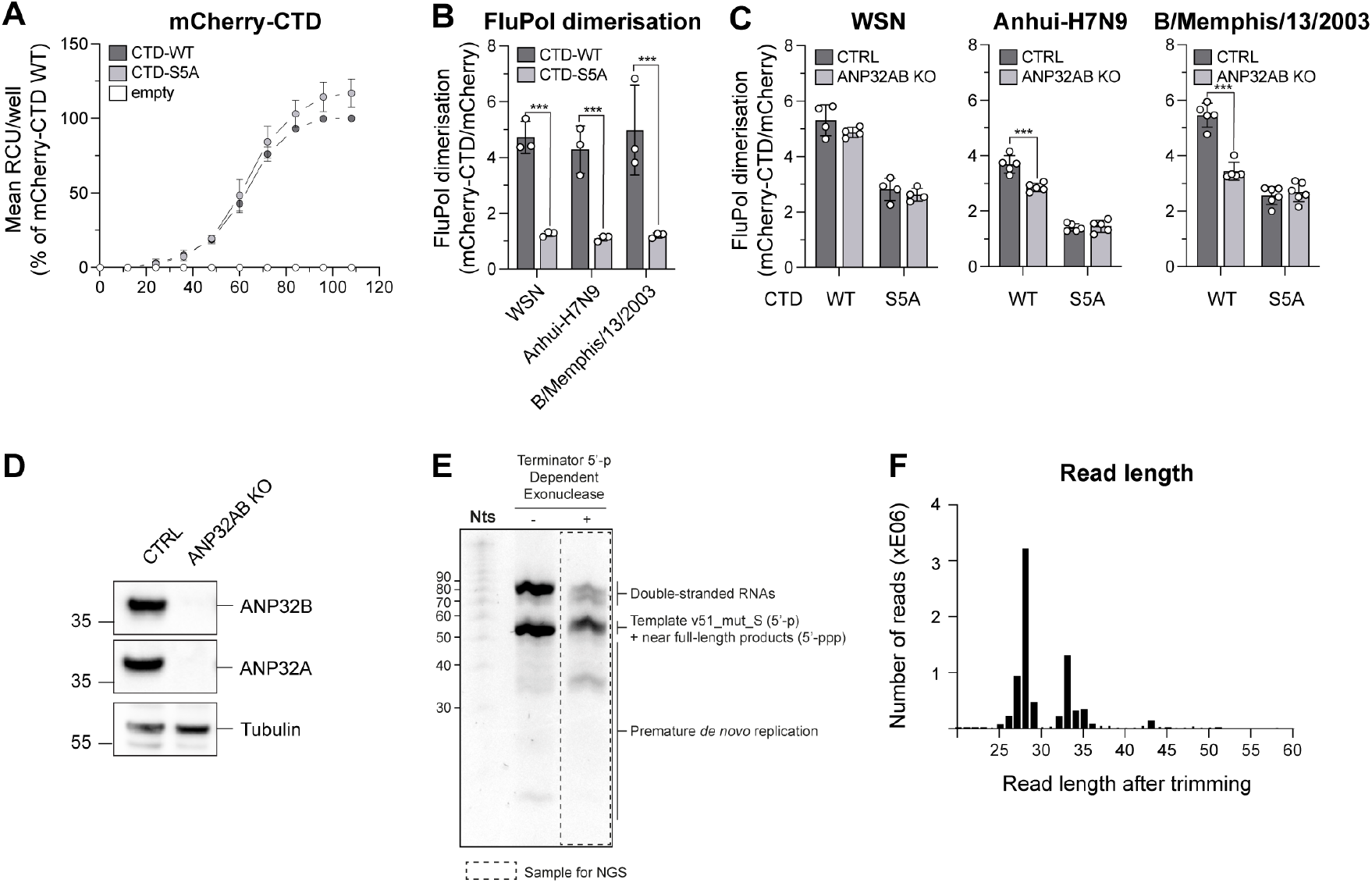
FluPol replication activity is enhanced in the presence of CTD and ANP32A. **(A)** HEK-293T cells were transiently transfected with expression plasmids encoding for mCherry-CTD-WT, mCherry-CTD-S5A or an empty plasmid control. Real-time monitoring of fluorescence was performed by the acquisition of 5 images with a 10X objective per well every 12 h up to 108 hpt. Mean fluorescence per well (RCU: Red Calibrated Units) is expressed as percentages of mCherry-CTD-WT at 108 hpt. **(B)** WSN, Anhui-H7N9, B/Memphis/13/2003 FluPol dimerisaftion was assessed in a split-luciferase-based complementation assay as described previously (Chen et al., 2019). Untagged mCherry or mCherry tagged to CTD-WT or CTD-S5A in which all serine 5 residues were replaced with alanines were co-expressed. Luminescence due to luciferase reconstitution was measured and the signals are represented as a percentage of untagged mCherry co-expression. ***p < 0.001 (two-way ANOVA; Sidak’s multiple comparisons test). **(C)** FluPol dimerisation levels of WSN, Anhui-H7N9 and B/Memphis/13/2003 were assessed as described in (B) either in HEK-293T control (CTRL) or in HEK-293T cells in which ANP32A and ANP32B were knocked out by CRISPR-Cas9 (ANP32AB KO). ***p < 0.001 (two-way ANOVA; Sidak’s multiple comparisons test). **(D)** HEK-293T CTRL and ANP32AB KO cell lysates were analysed by western blot using antibodies specific for ANP32A, ANP32B and tubulin. Uncropped gels are provided as a source data file. **(E)** NGS sample preparation of FluPol Zheijiang-H7N9 4M *de novo* replication products. Urea-PAGE gel stained with SYBR-Gold of the *de novo* reaction (Fig. 3 lane 8) before (-) and after (+) 5’-monophosphate (5’-p) RNA digestion using a terminator 5’-p-dependent exonuclease. The dotted rectangle corresponds to the sample sent for NGS. The decade molecular weight marker (Nts) is shown on the left side of the gel. The uncropped gel is provided as a source data file. **(F)** RNA-sequencing analysis of FluPol Zheijiang-H7N9 4M *de novo* replication products in the presence of pS5 CTD(6mer) and ANP32A. After trimming, the number of reads are plotted according to their lengths. NGS data are provided in source data table 1.

**Figure S6:**
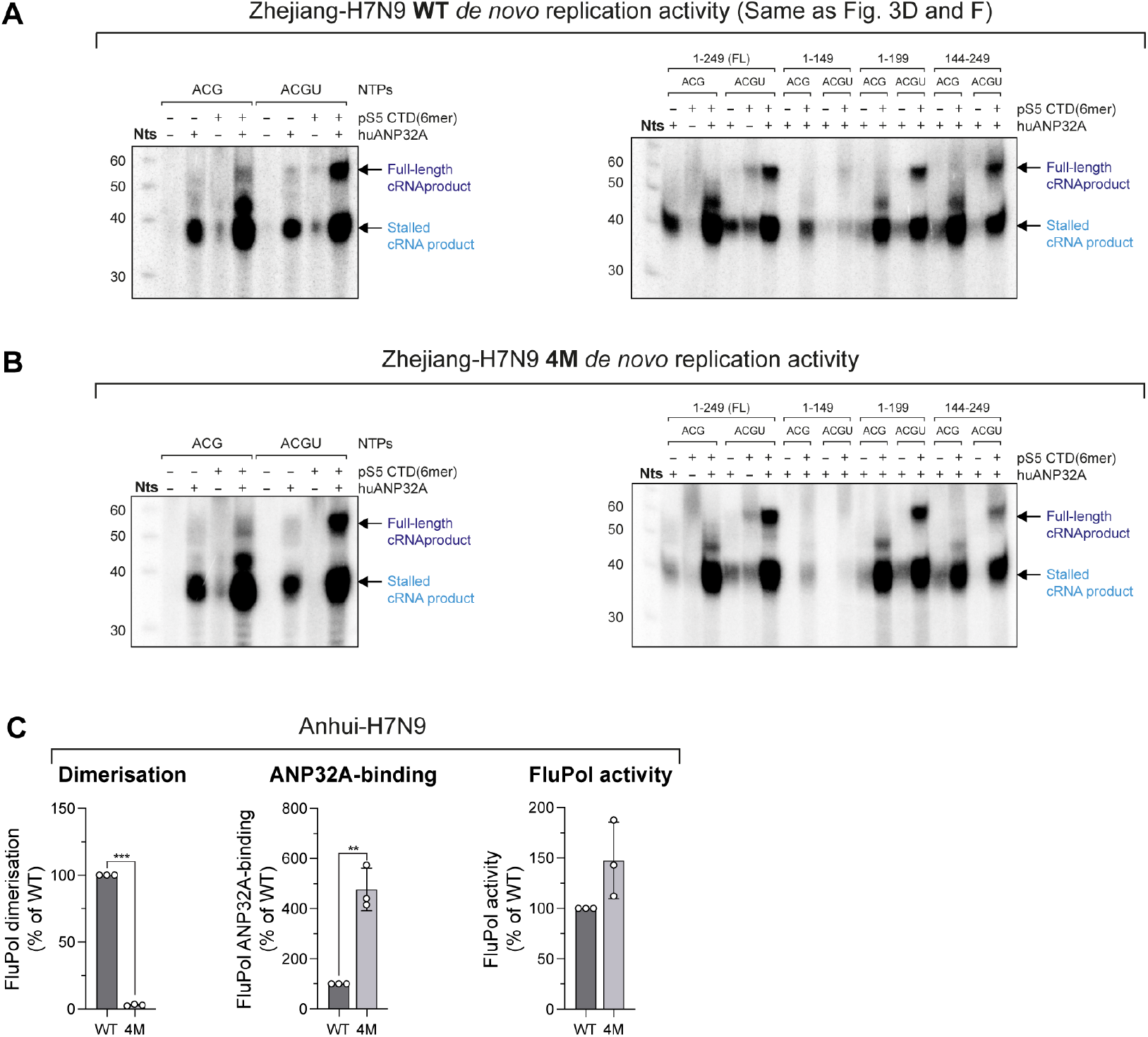
FluPol replication activity is enhanced in the presence of CTD and ANP32A. **(A-B)** *De novo* replication activity of (A) Zhejiang-H7N9 WT and (B) Zhejiang-H7N9 4M (PA E349K, PA R490I, PB1 K577G, and PB2-G74R) using the v51_mut_S template is enhanced in the presence of ANP32A and pS5 CTD(6mer). The assays are done in the presence of either 3 NTPs (AUG) or 4 NTPs (AUGC), with or without pS5 CTD(6mer) and ANP32A FL (left) or different ANP32A truncation mutants (right): ANP32A 1-249 (FL), 1-149, 1-199 and 144-249 as represented schematically in Fig. 3E. Full-length and stalled replication products are annotated on the right side of the gel and coloured as in Fig. 3C. The decade molecular weight marker (Nts) is shown on the left side of the gel. Uncropped gels are provided as a source data file. **(C)** Cell-based Anhui-H7N9 WT or 4M FluPol binding and activity assays. (left) FluPol dimerisation was assessed using a split-luciferase-based complementation assay as described previously (Chen et al., 2019). (middle) FluPol binding to ANP32A was assessed using a split-luciferase-based complementation assay. HEK-293T cells were co-transfected with expression plasmids for ANP32 tagged with one fragment of the *G. princeps* luciferase and FluPol tagged with the other fragment. Luminescence due to luciferase reconstitution was measured and the signals are represented as a percentage of PA WT. (right) FluPol activity was measured by vRNP reconstitution in HEK-293T cells, using a model vRNA encoding the Firefly luciferase. Luminescence was measured and normalised to a transfection control. The data are represented as a percentage of PA WT. **p < 0.002, ***p < 0.001 (one-way ANOVA; Dunnett’s multiple comparisons test).

**Figure S7:**
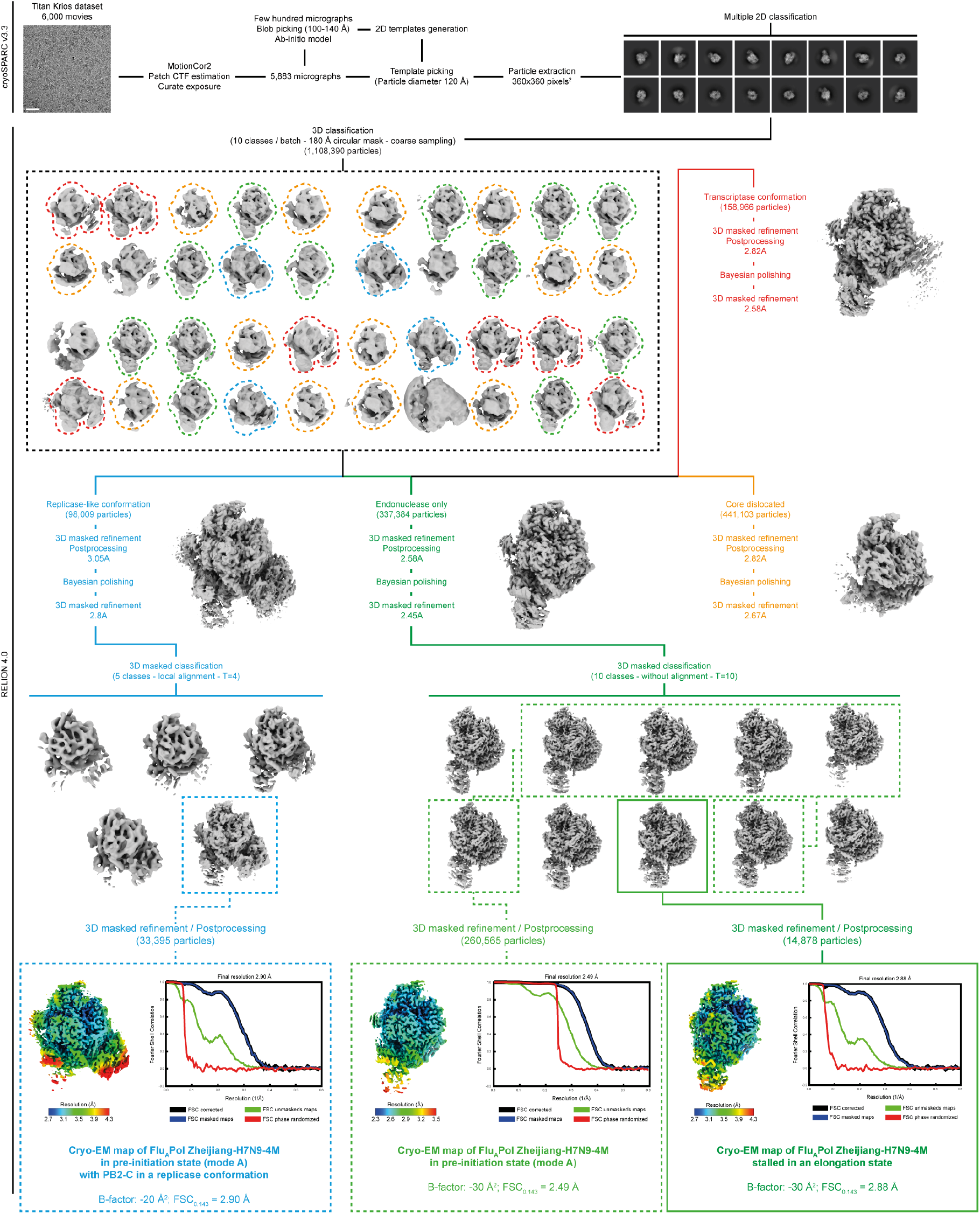
Image processing FluPol Zhejiang-H7N9-4M *de novo* replication reaction (Krios) Schematics of the image processing strategy used to obtain (i) the Cryo-EM map of FluPol Zhejiang-H7N9-4M in a pre-initiation state (mode A), with PB2-C in a replicase conformation, (ii) the cryo-EM map of FluPol Zhejiang-H7N9-4M in pre-initiation state (mode A) without PB2-C, and (iii) the cryo-EM map of FluPol Zhejiang-H7N9-4M stalled in an elongation state using UpNHpp. Further processing steps for the transcriptase conformation and dislocated FluPols are not shown. Representative cropped micrograph from the TEM Titan Krios dataset, 2D class averages, 3D class averages and intermediate structures are displayed. CTF stands for “Contrast Transfer Function”. The final number of particles, filtered EM maps according to the local resolution and corresponding Fourier Shell Correlation curves (FSC) are shown. Scale bar = 200 Å.

**Figure S8:**
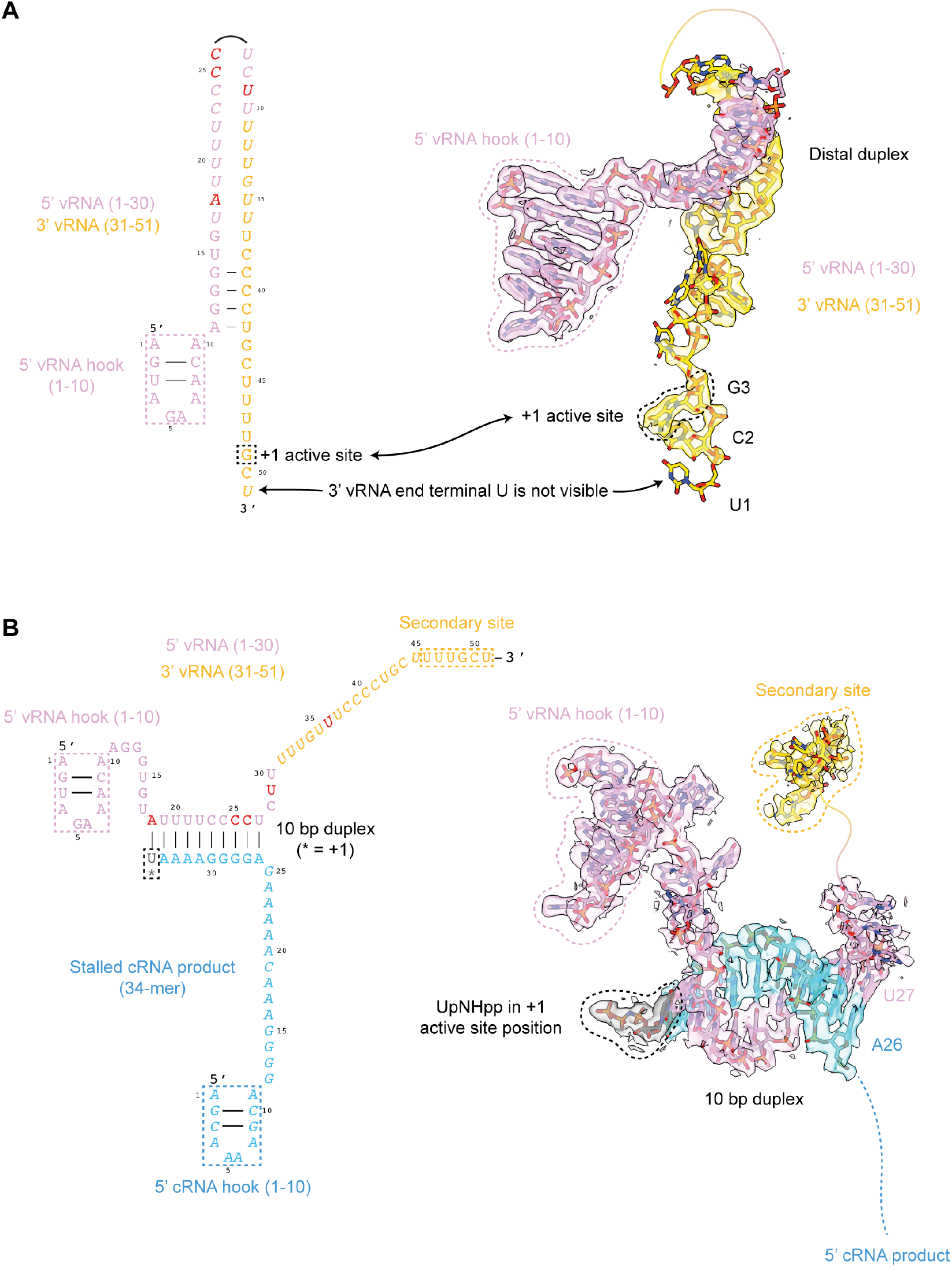
RNA conformation schematics, densities and models of the different states obtained. **(A)** (Left) Schematics of the v51_mut_S template conformation present in both (i) the cryo-EM map of FluPol Zhejiang-H7N9-4M in pre-initiation state (mode A) without PB2-C or (ii) with PB2-C in replicase conformation. The 5’ vRNA end (1-30) is coloured in pink. The 3’ vRNA end (31-51) is coloured in gold. Introduced mutations are coloured in red. Unseen nucleotides are in italics. The +1 active site position is indicated by a dotted rectangle. (Right) Coulomb potential of the v51_mut_S isolated from the cryo-EM map of FluPol Zhejiang-H7N9 4M in pre-initiation state (mode A) without PB2-C, at an overall resolution of 2.5 Å. 3’-U1 is displayed but not visible in the cryo-EM map. 3’-G3 is in the +1 active site position. **(B)** (Left) Schematics of the v51_mut_S template and the *de novo* cRNA replication product (34-mer) conformations present in the cryo-EM map of FluPol Zhejiang-H7N9-4M stalled in an elongation state using UpNHpp. The template is coloured as in A. The *de novo* cRNA product is coloured in blue. Unseen nucleotides are in italics. The +1 active site position is indicated. (Right) Coulomb potential of the v51_mut_S template and the *de novo* cRNA replication product (34-mer) isolated from the cryo-EM map of FluPol Zhejiang-H7N9-4M stalled in an elongation state using UpNHpp, at an overall resolution of 2.9 Å. The non-hydrolysable UpNHpp is in the +1 active site position, coloured in dark grey and circled by a dotted line. Flexible nucleotides are shown as dotted lines.

**Figure S9:**
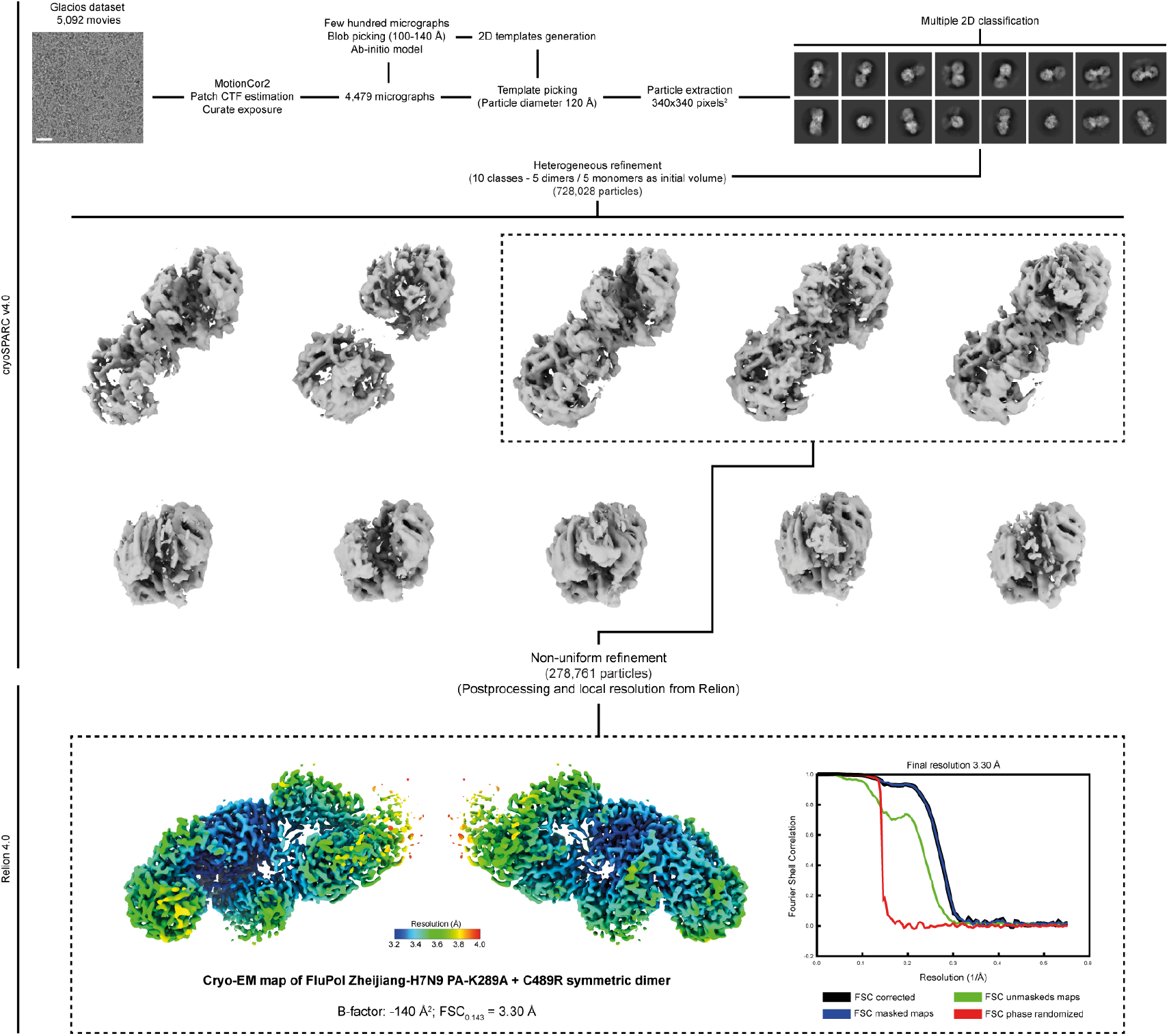
Image processing FluPol Zhejiang-H7N9 PA K289A+C489R (Glacios) Schematics of the image processing strategy used to obtain the cryo-EM map of FluPol Zhejiang-H7N9 PA K289A+C489R. Representative cropped micrograph from the TEM Glacios dataset, 2D class averages, 3D class averages and intermediate structures are displayed. CTF stands for “Contrast Transfer Function”. The final number of particles, filtered EM maps according to the local resolution and corresponding Fourier Shell Correlation curves (FSC) are shown. Scale bar = 200 Å.

**Figure S10:**
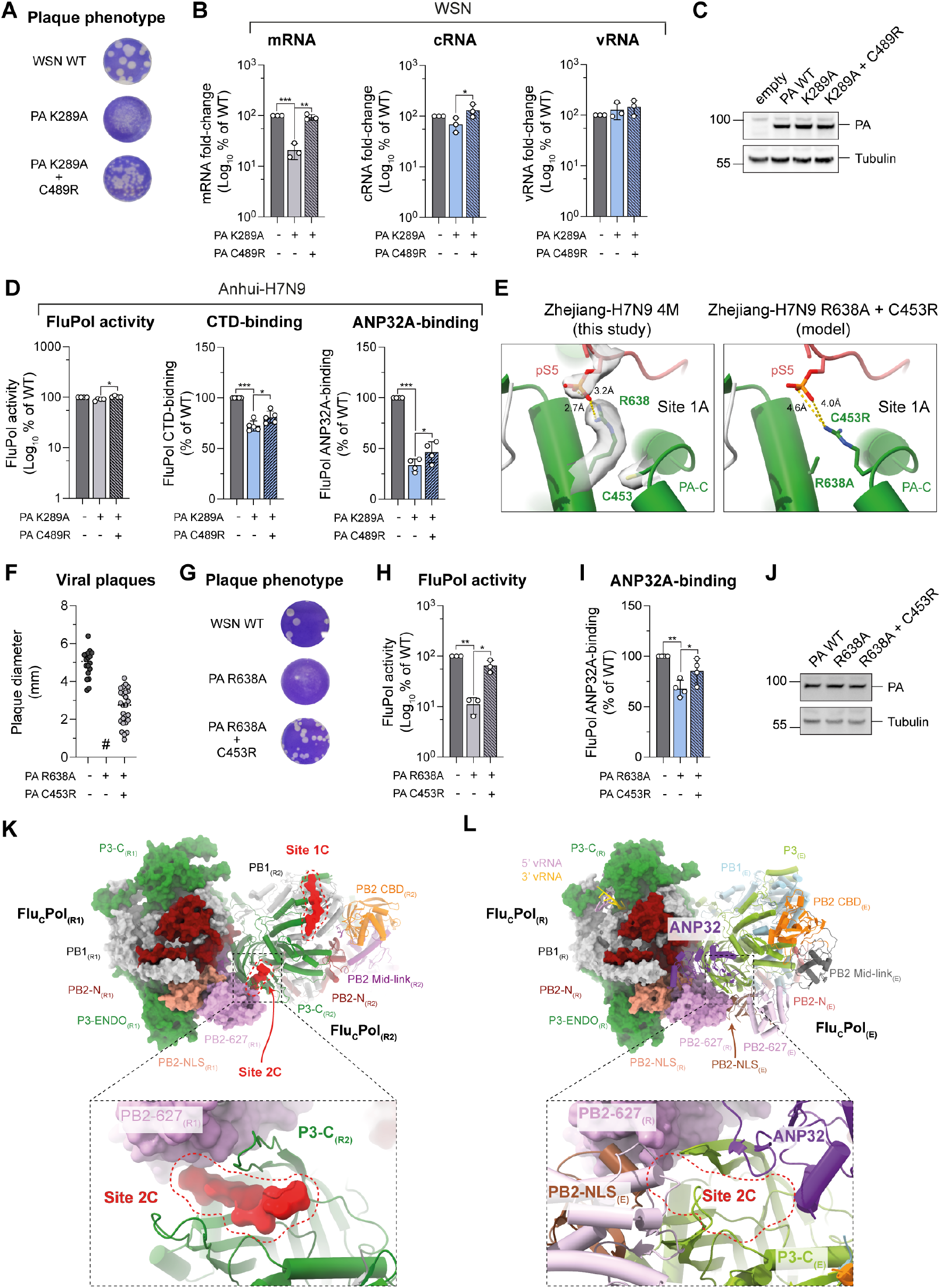
Restoration of CTD-binding enhances FluPol binding to ANP32A and FluPol replication activity. **(A)** Recombinant WSN mutant viruses were produced by reverse genetics. Representative plaque assays as quantified in Fig. 5D of mutant viruses harbouring PA K289A and the second-site mutation PA C489R are shown. The supernatants of two independently performed RG experiments were titrated on MDCK cells and stained by crystal violet. **(B)** WSN vRNPs with the PA K289A mutation or the double mutation PA K289A+C489R were reconstituted in HEK-293T cells by transient transfection using a plasmid encoding the NA vRNA segment. Steady-state levels of NA mRNA, cRNA and vRNA were quantified by strand-specific RT-qPCR (Kawakami et al., 2011) normalised to GAPDH by the 2^−ΔΔCT^ method (Livak and Schmittgen, 2001) and are represented as a percentage of PA WT. *p < 0.033, **p < 0.002, ***p < 0.001 (one-way ANOVA; Dunnett’s multiple comparisons test). **(C)** HEK-293T cells were co-transfected with expression plasmids for WSN PB1, PB2 and PA with the indicated mutation. Cell lysates were analysed by western blot using antibodies specific for PA and tubulin. Uncropped gels are provided as a source data file. **(D)** Anhui-H7N9 Flu_A_Pol with the PA K289A mutation and the second-site mutation PA C489R were characterised. (left) Anhui-H7N9 Flu_A_Pol activity was measured by vRNP reconstitution (PB2, PB1, PA-ΔX, NP) in HEK-293T cells, using a model vRNA encoding the Firefly luciferase. Luminescence was measured and normalised to a transfection control. The data are represented as a percentage of PA WT. (middle) Anhui-H7N9 Flu_A_Pol binding to the CTD was assessed using a split-luciferase-based complementation assay. HEK-293T cells were co-transfected with expression plasmids for the CTD tagged with one fragment of the *G. princeps* luciferase and Flu_A_Pol tagged with the other fragment. Luminescence due to luciferase reconstitution was measured and the data are represented as a percentage of PA WT. (right) ANP32A was tagged with one fragment of the *G. princeps* luciferase and binding to WSN Flu_A_Pol was determined as for Flu_A_Pol l-binding to the CTD. *p < 0.033, ***p < 0.001 (one-way ANOVA; Dunnett’s multiple comparisons test). **(E)** (left) Cartoon representation of the pS5 CTD bound to Flu_A_Pol Zhejiang-H7N9 z4M (replicase-like conformation) in CTD-binding site 1A. The PA subunit is coloured in green and the pS5 CTD is coloured in red. PA C453 and PA R638 residues are displayed. Putative hydrogen bonds between PA R638 and pS5 are drawn as yellow dashed lines and corresponding distances are indicated. The Coulomb potential map of PA C453, PA R638, and pS5 is shown. (right) Model derived from Flu_A_Pol Zhejiang-H7N9 4M structure bearing PA C453R and PA R638A mutations. The PA subunit is coloured in green and the pS5 CTD is coloured in red. PA C453R and PA R638A residues are displayed. Hypothetical distances between PA C453R and pS5 are shown. **(F-I)** Phenotypes associated with the WSN Flu_A_Pol PA R638A mutation and the second-site mutation PA C453R were characterised. (F) Plaque phenotype of recombinant WSN mutant viruses produced by reverse genetics. RG supernatants were titrated on MDCK cells and stained by crystal violet. Plaque diameters (mm) were measured and each dot represents one viral plaque. Representative plaque assays are shown in (G) (#) not measurable pinhead-sized plaques. (H) WSN Flu_A_Pol activity was measured by vRNP reconstitution in HEK-293T cells, using a model vRNA encoding the Firefly luciferase. Luminescence was measured and normalised to a transfection control. The data are represented as a percentage of PA WT. (I) WSN Flu_A_Pol binding to the ANP32A was assessed using a split-luciferase-based complementation assay. HEK-293T cells were co-transfected with expression plasmids for the CTD tagged with one fragment of the *G. princeps* luciferase and Flu_A_Pol tagged with the other fragment. Luminescence due to luciferase reconstitution was measured and the data are represented as a percentage of PA WT. *p < 0.033, **p < 0.002 (one-way ANOVA; Dunnett’s multiple comparisons test). **(J)** HEK-293T cells were co-transfected with expression plasmids for WSN PB1, PB2 and PA with the indicated mutation. Cell lysates were analysed by western blot using antibodies specific for PA and tubulin. V **(K)** Flu_C_Pol asymmetric dimer bound to pS5 CTD (PDB: 6F5P, (Serna Martin et al., 2018). Both Flu_C_Pols are in replicase conformation. The first Flu_C_Pol_R1_ (Replicase(1)) is displayed as surface. The second Flu_C_Pol_R2_ (Replicase(2)) is displayed as cartoon. P3 subunit is coloured in green, PB1 subunit in light grey, PB2-N in dark red, PB2 Mid-link in purple, PB2 CBD in orange, PB2 627 in pink, PB2 NLS in beige. Both pS5 CTD binding sites (1C, 2C) are circled in red. Each bound pS5 CTD peptides are displayed as surface and coloured in red. Close-up view on pS5 CTD bound to the site 2C is shown below. **(L)** Flu_C_Pol asymmetric dimer bound to ANP32A (PDB: 6XZG, (Carrique et al., 2020)). The first Flu_C_Pol is in a replicase conformation (Replicase_(R)_), displayed and coloured as in K. The 5’ vRNA end is coloured in pink and the 3’ vRNA end is coloured in gold. The second Flu_C_Pol is in an encapsidase conformation (Encapsidase_(E)_), displayed as cartoon. P3_E_ subunit is colored in light green, PB1_E_ subunit in light blue, PB2-N_(E)_ in salmon, PB2 Mid-link_(E)_ in dark grey, PB2 CBD_(E)_ in orange, PB2 627_(E)_ in light pink, PB2 NLS_(E)_ in brown. ANP32A is displayed as cartoon and coloured in purple. Close-up view on the pS5 CTD in binding site 2C is shown.

